# Resistance to Naïve and Formative Pluripotency Conversion in RSeT Human Embryonic Stem Cells

**DOI:** 10.1101/2024.02.16.580778

**Authors:** Kevin G. Chen, Kory R. Johnson, Kyeyoon Park, Dragan Maric, Forest Yang, Wen Fang Liu, Yang C. Fann, Barbara S. Mallon, Pamela G. Robey

## Abstract

One of the most important properties of human embryonic stem cells (hESCs) is related to their primed and naïve pluripotent states. Our previous meta-analysis indicates the existence of heterogeneous pluripotent states derived from diverse naïve protocols. In this study, we have characterized a commercial medium (RSeT)-based pluripotent state under various growth conditions. Notably, RSeT hESCs can circumvent hypoxic growth conditions as required by naïve hESCs, in which some RSeT cells (e.g., H1 cells) exhibit much lower single cell plating efficiency, having altered or much retarded cell growth under both normoxia and hypoxia. Evidently, hPSCs lack many transcriptomic hallmarks of naïve and formative pluripotency (a phase between naive and primed states). Integrative transcriptome analysis suggests our primed and RSeT hESCs are close to the early stage of post-implantation embryos, similar to the previously reported primary hESCs and early hESC cultures. Moreover, RSeT hESCs did not express naïve surface markers such as CD75, SUSD2, and CD130 at a significant level. Biochemically, RSeT hESCs exhibit a differential dependency of FGF2 and co-independency of both Janus kinase (JAK) and TGFβ signaling in a cell-line-specific manner. Thus, RSeT hESCs represent a previously unrecognized pluripotent state downstream of formative pluripotency. Our data suggest that human naïve pluripotent potentials may be restricted in RSeT medium. Hence, this study provides new insights into pluripotent state transitions *in vitro*.

## INTRODUCTION

Naïve or ground pluripotent states were initially proposed by Smith and colleagues based on studies in mouse embryonic stem cells (mESCs) from pre-implantation mouse embryos (Nichols and Smith, 2009; Ying et al., 2008) and on studies of lineage-primed epiblast stem cells (EpiSCs) derived from the post-implantation mouse epiblast (Brons et al., 2007; Tesar et al., 2007). The establishment of human naïve pluripotency is believed to have significant impacts on understanding early human embryonic development, facilitating single human pluripotent stem cell (hPSC) growth *in vitro*, genetic manipulation, disease-modeling, optimizing interspecies chimerism, and drug discovery [reviewed in (Dodsworth et al., 2015; Dong et al., 2019; Hanna et al., 2010; Weinberger et al., 2016; Wray et al., 2010; Wu et al., 2017; Zimmerlin et al., 2017)]. In the past decade, numerous groups reported the conversion of primed hPSCs to naïve hPSCs or direct derivation of them from the human inner cell mass (ICM) (Chan et al., 2013; Collier et al., 2017; Gafni et al., 2013; Guo et al., 2016; Kilens et al., 2018; Liu et al., 2017; Takashima et al., 2014; Theunissen et al., 2016; Ware et al., 2014; Zimmerlin et al., 2016). Despite the enthusiasm of deriving naïve hPSCs with diverse growth conditions, there are increasingly intense efforts to derive naïve hPSCs based on the use of the commercial medium RSeT (www.stemcell.com) (Collier et al., 2017; Kilens et al., 2018; Liu et al., 2017; Szczerbinska et al., 2019). Our meta-analysis indicated that RSeT-based hPSCs have a closer transcriptomic similarity to primed and formative hPSCs than other “naïve” counterparts derived from different laboratories (Johnson et al., 2021). Likely, RSeT hPSCs provide an important clue to unraveling human naïve and formative pluripotency in development and in cell culture platforms. However, the cellular properties and pluripotent states in RSeT hPSCs under different growth conditions are not well defined or characterized.

In this study, we report the conversion and more detailed characterization of RSeT hESCs, including transcriptomic analysis of primed and naïve gene expression, diagnostic cell surface marker expression, and the response to core signaling molecules of both primed and naïve pluripotency. We aim to provide a systemic evaluation of RSeT conditions and resolve some experimental and conceptual inconsistencies concerning human naïve and formative pluripotency (i.e., a transition phase between the naïve and primed states of epiblast) (Smith 2017). Our data reveal that RSeT medium does not effectively support the conversion of primed hESCs to a naïve state. The converted RSeT hESCs have a distinct and previously unrecognized pluripotent state between the formative and primed states, displaying insensitivity to oxygenic tension alterations, altered response to core signaling molecules, and significantly different cell growth potentials.

## RESULTS

### Conversion of primed hESC lines to RSeT states without the hypoxic growth requirements for naïve hESCs

RSeT medium is based on naïve human stem cell medium (NHSM) (Gafni et al., 2013), but with a different formulation (www.stemcell.com). Our investigation revealed that RSeT-converted hPSCs exhibit a transcriptomic profile akin to primed hPSCs (Johnson et al., 2021), although it was not previously recognized (refer to Figure 1A). To fully characterize RSeT hPSCs, we converted three primed hESC lines (H1, H7, and H9) under RSeT growth conditions (Figure 1B), with the initial use of the ROCK inhibitor Y-27632 (ROCKi) to enhance single-cell plating efficiency (SCPE).

**Figure 1.**
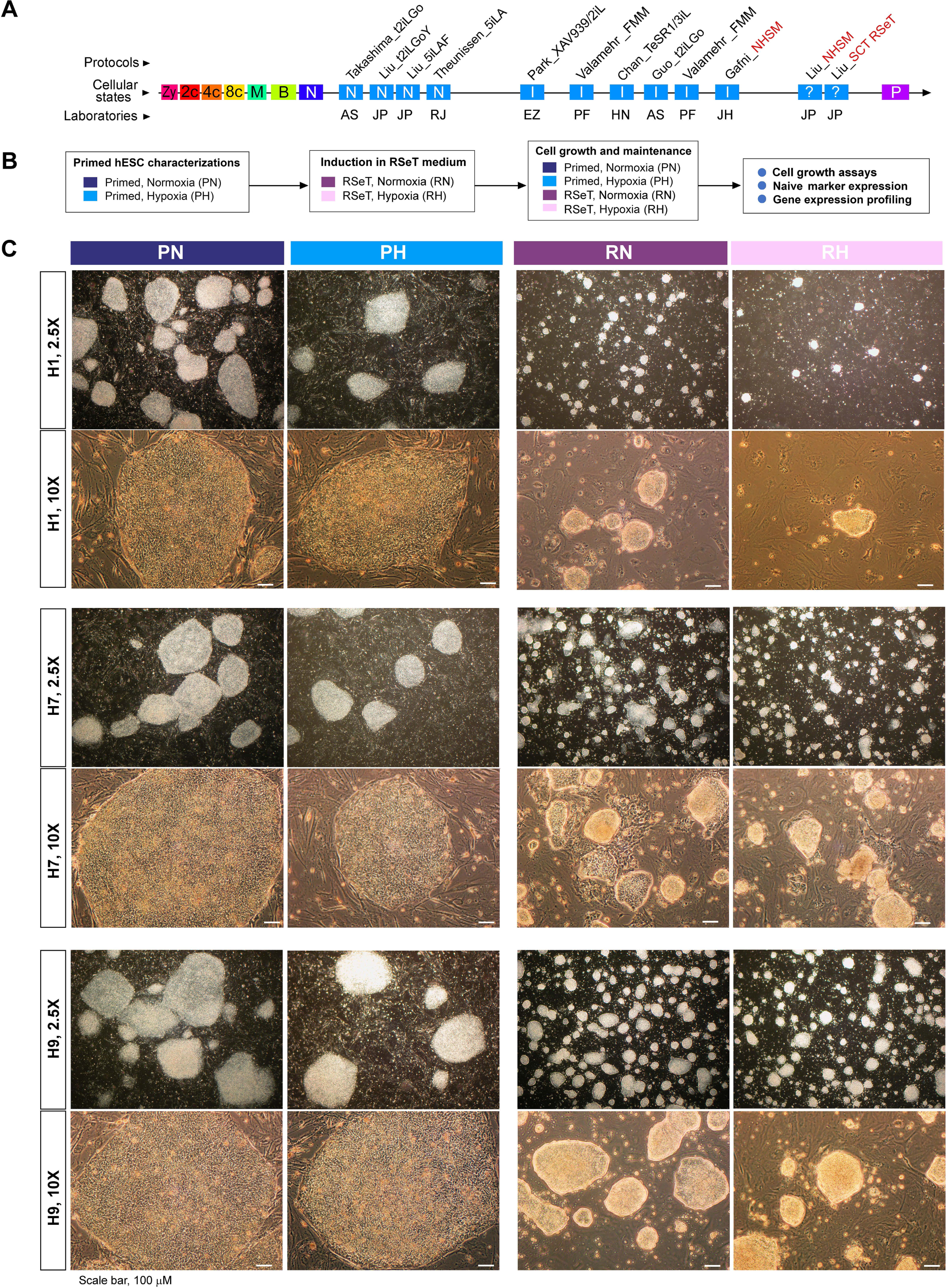
RSeT conversion of human embryonic stem cells (hESCs) under normoxia and hypoxia. (**A**) Linear presentation of diverse hPSC pluripotent states determined by meta-analysis (Johnson et al., 2021), with current naïve conversion or reprogramming protocols (labeled in italicized characters with first authors’ names), t2iLGo, t2iLGoY, 5iLA, 5iLAF, XAV939/2iL, FMM, TeSR1/3iL, NHSM, and RSeT, from the laboratories of AS (Austin Smith), EZ (Elias Zambidis), HN (Huck-Hui Ng), JH (Jacob Hanna), JP (Jose Polo), PF (Peter Flynn), and RJ (Rudolf Jaenisch). The hPSCs (in red-font characters) are related to RSeT protocols (SCT RSeT and NHSM). (**B**) Schema of hESC lines (H1, H7, and H9) treated under different primed and RSeT conditions as indicated. (**C**) Phase images of primed and RSeT hESCs grown on MEFs: PN, primed growth under normoxia; PH, primed growth under hypoxia; RN, RSeT growth under normoxia; and RH, RSeT growth under hypoxia. Additional abbreviations: 2c, 4c, and 8c: 2-cell, 4-cell, and 8 cell embryos, respectively; B, blastocyst; I, intermediate transitions between the naive and primed states; M, morulae; N, cells with the naïve pluripotent state; P, primed pluripotent states; and Zy, zygote.

The converted RSeT hESCs displayed a slow growth pattern with morphologically small domed-shaped colonies as described for naïve hPSCs (Figure 1C). However, we found that all three hESC lines behaved differently under the RSeT conditions. In general, H9 cells were the easiest to be induced into and maintained, as domed colonies (Figure 1C, bottom panel), followed by H7 cells (Figure 1C, middle panel). H1 cells displayed intractability to being induced and maintained as domed colonies under these conditions (Figure 1C, top panel). Upon removal of ROCKi after 3 passages, H1 cells exhibited extremely poor growth that was even exacerbated under hypoxia (i.e., 3% O_2_). For example, H1 cells exhibited a 6.7-fold reduction in colony formation at passage 10 under RSeT and hypoxic growth condition (RH) (Figure 1C, top panel). Interestingly, RSeT hESCs typically propagate well in normoxic conditions compared with the same cells under hypoxia (Figure 1C, RN and RH columns). However, H7 colonies grown in RSeT under normoxia (RN) showed some heterogeneous morphologies (Figure 1C, middle panel). Nonetheless, the data suggest that RSeT hESCs can bypass the hypoxic growth requirement for naïve hPSCs.

### RSeT hESCs transcriptomically similar to their primed counterparts

Globally, RSeT hESCs have a transcriptome similar to their primed hESCs counterparts based on microarray profiling of gene expression in this study (Figure 2A, Figure S1). Noticeably, both primed and RSeT transcriptomes comprise the majority (> 65%) of transcripts with low to moderate mRNA expression. Only a small portion of the mRNA transcripts were highly expressed in RSeT cells (Figure 2A). We further utilized principal component analysis (PCA) to define the differences between individual cell lines. PCA transforms a large number of variables into a smaller number of orthogonal variables (e.g., principal components 1, 2, and 3, abbreviated as PC1, PC2, and PC3, respectively). Hence, 96.1% of gene expression variations, in which PC1 and PC2 constitute 88.5% and 7.6% variability respectively, might be used to depict the differences between primed and RSeT states (Figure 2B). Moreover, H9 cells showed a similar transcriptome to H1 cells under primed growth conditions, but closer to H7 than H1 under RSeT conditions (Figure 2A; 2B, left panel). Interestingly, only a small percentage (i.e., 2.4%) of gene expression variations was responsible for the response to normoxic and hypoxic growth (Figure 2B, right panel), which may be associated with the enhanced growth capacity of RSeT hESCs under normoxia as described above.

**Figure 2.**
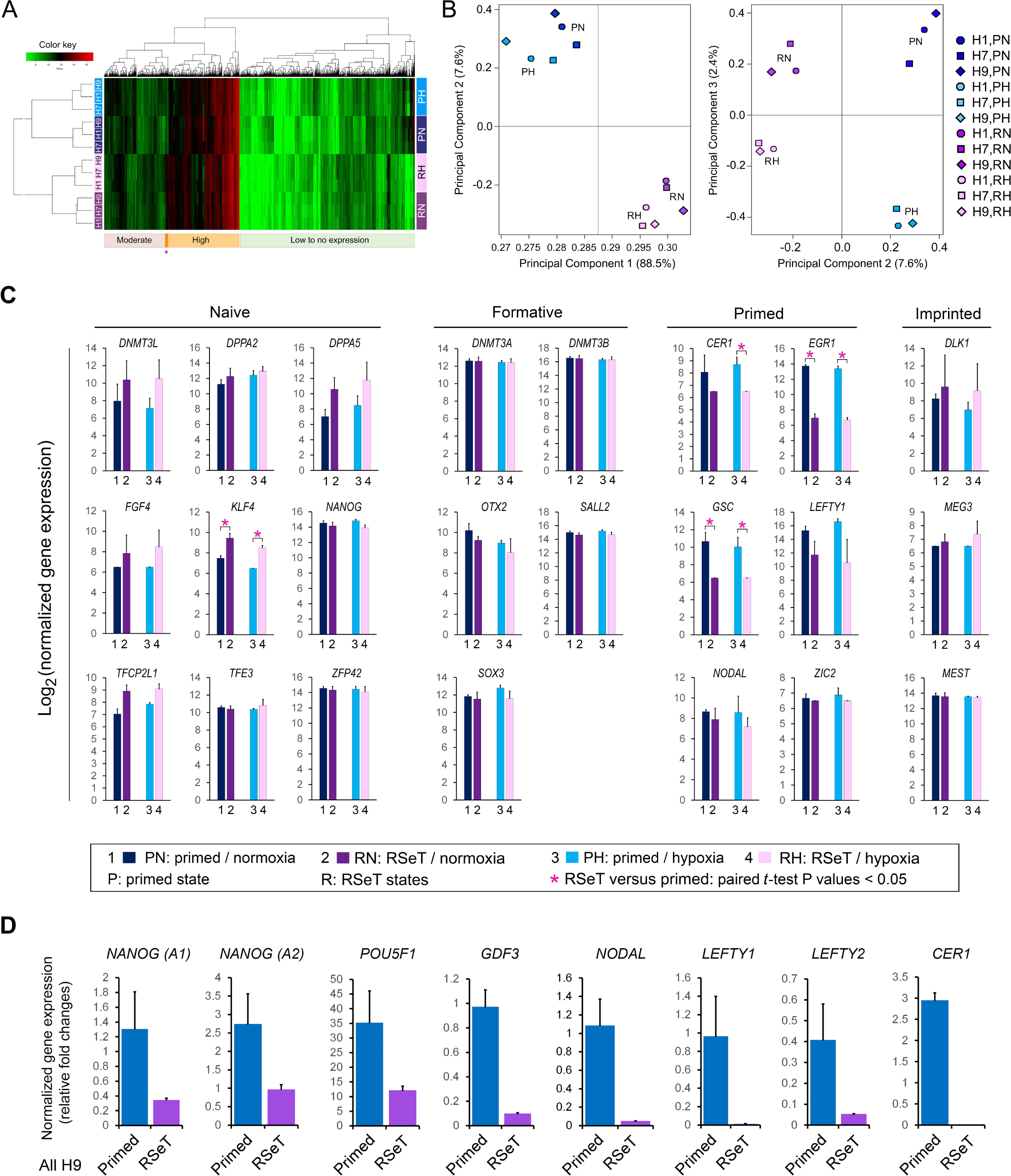
Genome-wide transcriptomic analysis of RSeT hESCs. **(A)** cDNA microarray heatmap of mRNA expression in RSeT hESCs. (**B**) Covariance-based principal component analysis (PCA) of the variability of gene expression as shown in Figure 2A. (**C**) Representative naïve, formative, primed, and imprinted gene markers from microarray analysis. (**D**) Quantitative real-time PCR (qPCR) confirmation of gene expression patterns based on microarray.

### RSeT hESCs lack naïve and formative gene markers

To define naïve and formative gene expression, we implemented a supervised analysis of a 112-gene list (Table S1), derived from previous mouse and human pluripotent stem cell reports or reviews (Brons et al., 2007; Chan et al., 2013; Collier et al., 2017; Gafni et al., 2013; Liao et al., 2015; Mallon et al., 2013; Pera and Rossant, 2021; Smith, 2017; Takashima et al., 2014; Tesar et al., 2007; Theunissen et al., 2016; Valamehr et al., 2014; Weinberger et al., 2016), which defines tangible primed and naïve transcriptomes. Briefly, this list includes the ICM and other developmental markers that are not expressed in the implantation epiblast (n = 5), naïve or epiblast-specific genes (n = 60), formative (between pre-implantation and the E6.5 epiblast in mouse) genes (n = 5), primed or metastable lineage specifier genes (n = 37), and other X-linked or imprinted gene expression (n = 5).

Overall, there are approximately 45% of genes within the 112-gene cluster that had altered expression when converted to RSeT cells. Some of the most notable human naïve signature genes (e.g., *DNMT3L*, *DPPA2*, *DPPA5*, *TFCP2L1*, *XIST*, and *ZFP42)* remained unchanged by microarray assays (Figure 2C, left panel; Table S1). Also, there was only 4.3% of genes altered between primed and RSeT states (Table S1), which is consistent with genome-wide microarray analysis showing the limited alterations of gene expression between the two states of hESCs (Figures 2A, 2B).

Notably, only a few naïve pluripotency genes (e.g., *KLF4*) were upregulated (Figure 2C, Tables S1). Interestingly, *NANOG* was downregulated in RSeT cells in both RSeT normoxia (RN) and RSeT hypoxia (RH) conditions (Figure 2C, left panel), which was confirmed by quantitative real-time PCR (qPCR) analysis using two different *NANOG* probes (Figure 2D, n = 6 cell lines, *P* < 0.05). *POU5F1*, another important pluripotency marker gene, was also significantly suppressed in RSeT hESCs as determined by qPCR (Figure 2D, n = 6, *P* < 0.05). Moreover, the expression of *GDF3* and *NODAL,* which were highly expressed in the naïve epiblast of human embryos (Blakeley et al., 2015; Stirparo et al., 2018), was also significantly reduced in RSeT hESCs (Figure 2D, n = 6, *P* < 0.05). These results are clearly opposite to that which was observed in reported naïve hPSCs.

Moreover, several formative pluripotency associated genes (e.g., *DNMT3A*, *DNMT3B, OTX2, SALL2, and SOX3*) and imprinted gene clusters (e.g., *DLK1*, *MEG3*, and *MEST*) had no significant alterations (Figure 2C, middle and right panels; Table S1). Other down-regulated primed genes include *CER1, EGR1, GSC, LEFTY1, NODAL, and ZIC2* (Figure 2C, Table S1). Some of which (e.g., *CER1*, *LEFTY1*, and *NODAL*) were also confirmed by our qPCR analysis (Figure 2D). Taken together, RSeT hESCs do not vary significantly in terms of their global gene signatures when compared with their primed parental cells. Specifically, RSeT hESCs lack human naïve and formative transcriptomic hallmarks as described in other human naive hPSC models, which makes it hard to ascertain the developmental stage of RSeT hESCs.

### Meta-analysis revealed a unique transcriptome and plausible developmental stage of RSeT hESCs

We utilized a comparative meta-analysis based on both PCA and *t*-distributed stochastic neighbor embedding (*t*-SNE) to resolve the above issue (Figure 3; Figures S2 and S3). This analysis revealed that RSeT hESCs had a minimal transcriptomic difference compared with their primed counterparts (Figures 3A-C), which were in contrast to some established naïve hPSCs, particularly to those derived by t2iLGo and 5iLAF protocols (Takashima et al., 2014; Theunissen et al., 2016) (Figures 3A-C).

**Figure 3.**
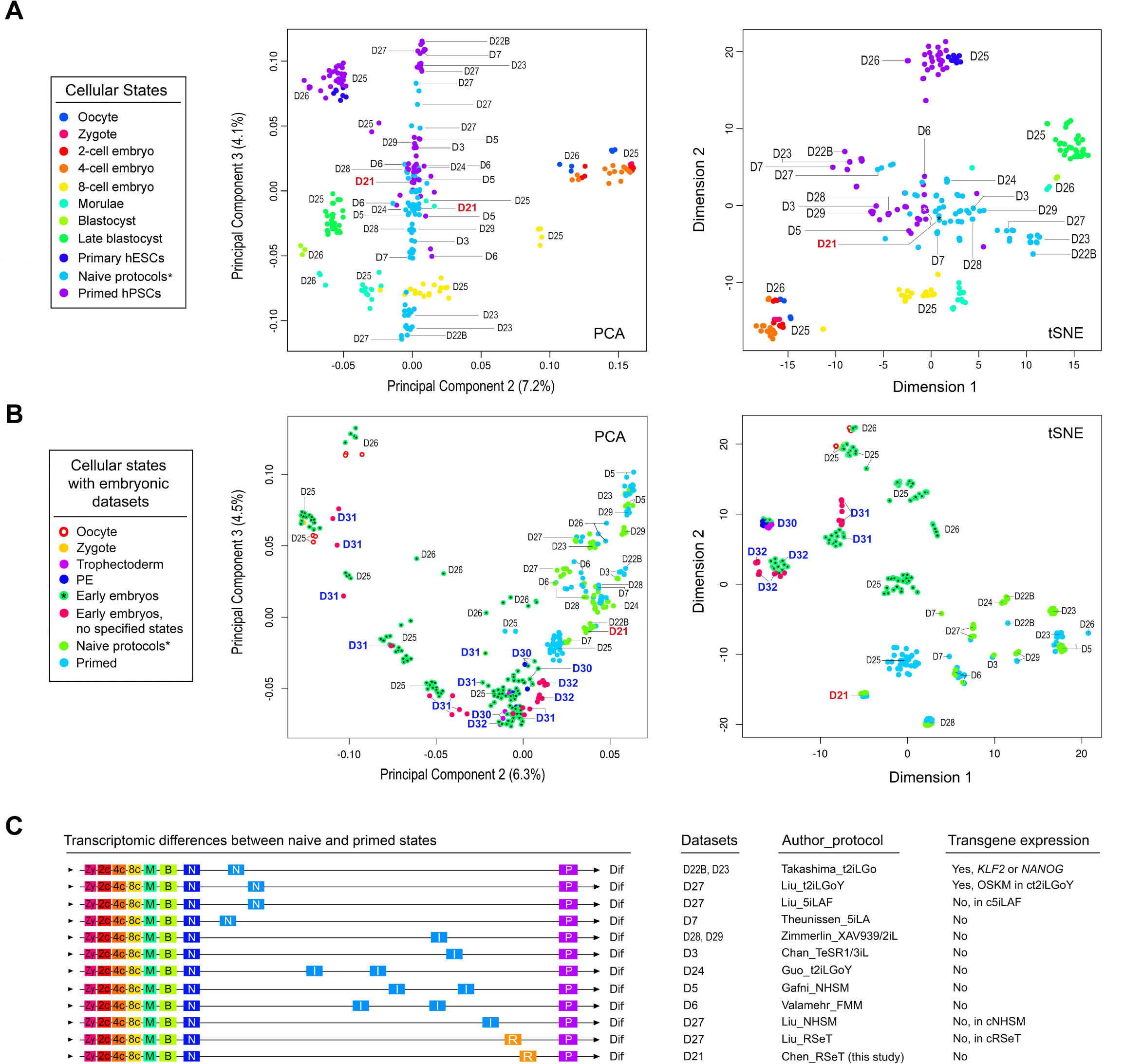
Genome-wide transcriptomic analysis of RSeT hESCs and meta-analysis of human naïve pluripotency. (**A**) Comparative meta-analysis of hPSC datasets generated from various protocols by Pearson correlation-based PCA and *t*-distributed stochastic neighbor embedding (*t*-SNE), which depict cellular and pluripotent states (color-coded dots) with dataset identification numbers as labeled (D3 to D29). The datasets used for comparison in this study were designated based on the name of first authors of published reports as detailed previously (Johnson et al., 2021). (**B**) PCA and *t*-SNE of detailed pluripotent states and developmental stages with the addition of multiple early primate embryo datasets (D30 to D32). (**C**) Mapping cellular and pluripotent states based on the Euclidean distance of gene-wide transcriptomic datasets. Abbreviations for cellular states and human naïve protocols (including the expression of transgenes) are indicated, which are also detailed in Materials and Methods and in the legend to Figure 1A. Additional abbreviations: Dif, lineage differentiation; I, intermediate transitions between the naïve and primed states; N, cells with naive pluripotent states; OKSM, transgene expression of the Yamanaka factors OCT4, KLF4, SOX2, and c-MYC; P, primed pluripotent states; R, hPSCs with a distinct pluripotent state in RSeT medium.

To directly relate cultured hPSCs to a developmental stage represents a significant challenge because of the presence of substantial batch effects between cultured hESCs and embryo datasets. However, we were able to maximally reduce the inter-laboratory and cell-culture batch effects (Johnson et al., 2021). This was further confirmed by including more primate embryo datasets, where all the primate embryo data are clustered closer together (Figure 3B, S3). Notably, the primed hESCs maintained in our laboratory are significantly different from their primed counterparts from several different laboratories (Figure 3B). Thus, we can associate our primed and RSeT hESCs with the early stage of post-implantation embryos, similar to the previously reported primary hESCs (at passage 0) and early hESC cultures at passage 10 (Figures 3B, S3) (Yan et al., 2013). These primary hESCs were derived from the ICM of blastocysts at day 6 (E6) with the least cell-culture-related changes (Yan et al., 2013).

Consequently, the findings of this analysis suggest that both primed and RSeT hESCs cultivated in our laboratory exhibit a similar transcriptomic state, and further indicate that these cells are not significantly affected by in vitro cell culture artifacts. Nonetheless, this observation emphasizes the presence of heterogeneous primed states under various cell culture conditions. It is worth noting that the primed hESCs grown in our laboratory may not be completely primed due to their resemblance to primary hESCs (Yan et al., 2013) (Figure 3B, S3), potentially harboring some unprimed naïve hESCs.

### Feeder-supported hESCs are not fully primed and heterogeneously express **naïve**-specific surface markers

We initially showed that RSeT hESCs highly expressed several primed pluripotency surface markers such as SSEA-4, Tra-1-60, and Tra-1-81 (Figure 4A). To verify the primed state of hESCs, we also examined the primed hESCs cells with several well-characterized primed-specific markers (e.g., CD24, CD57, and CD90) and naïve-specific markers (e.g., CD75, SUSD2, and CD130) (Bredenkamp et al., 2019; Collier et al., 2017) by flow cytometric analysis of Hoechst 33258-gated single cells. Surprisingly, the low oxygen level led to the partial expression of CD57 (39% in H1 and H7, and 45% in H9 cells), CD24 (58% in H1, 29% in H7, and 21% in H9 cells), and CD90 (41% in H1, 35% in H7, and 51% in H9 cells) in primed hESCs (Figures 4B-D, 5A1).

**Figure 4.**
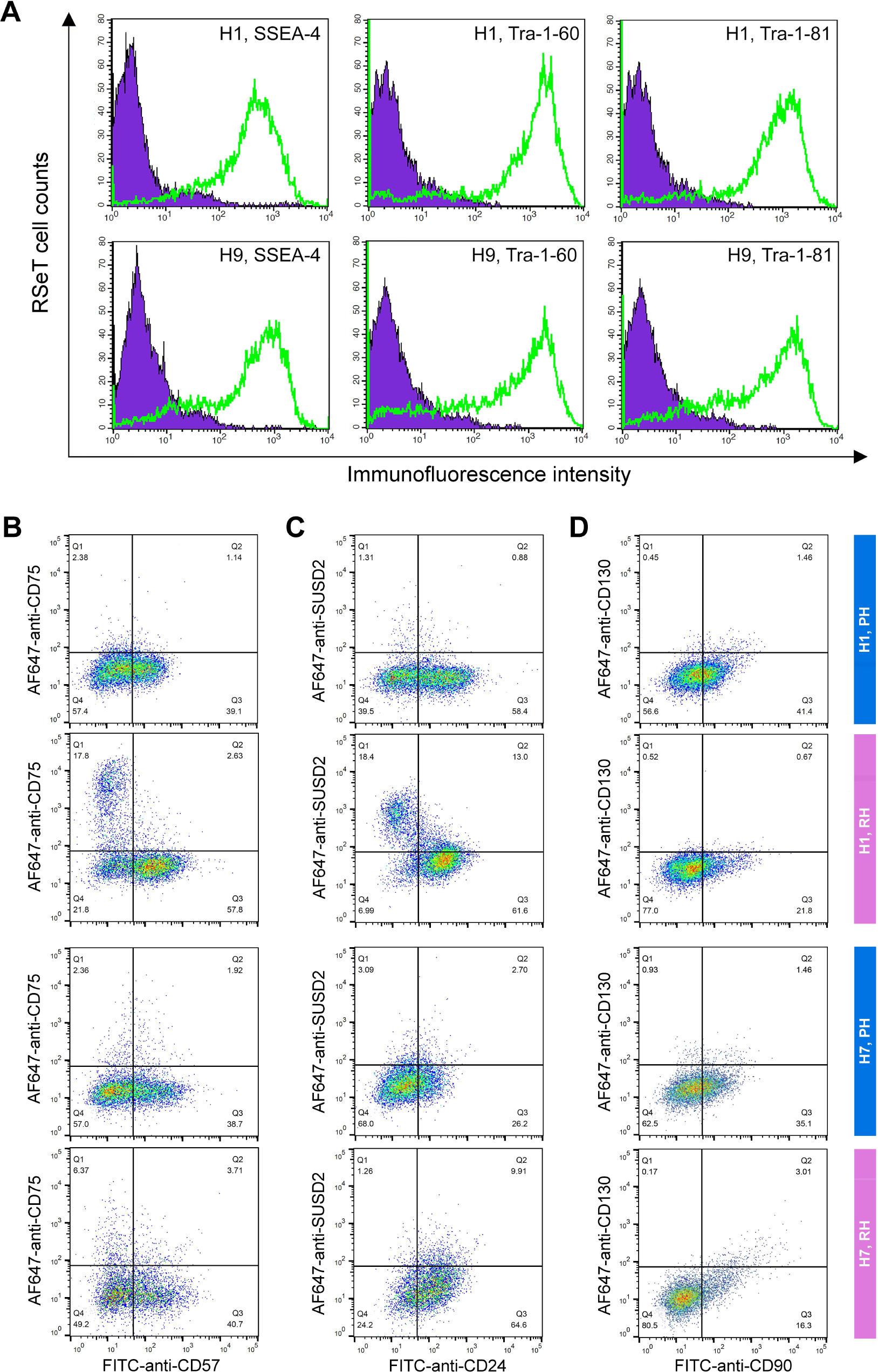
Flow cytometric analysis of human pluripotent stem cell surface marker expression. **(A)** Representative analysis of the expression of the human pluripotent stem cell surface markers SSEA-4, Tra-1-60, and Tra-1-81 in RSeT H1 and H9 cells. (**B to D**) Expression of the primed-specific markers (CD57, CD24, and CD90) as well as the naïve-specific surface markers (CD75, SUSD2, and CD130) in Hoechst 33258-gated H1 and H7 single cells. PH, primed hESCs grown under hypoxia; RH, RSeT-converted hESC grown under hypoxia.

Moreover, we found a very low percentage (0.5 to 3.1%) of primed H1 and H7 hESCs expressing naïve-specific markers CD75, SUSD2, and CD130 (Figures 4B-D). Unexpectedly, 14% of primed H9 cells express the naïve-specific marker SUSD2 (Figure 5A1). These data suggest that hESCs maintained on mouse embryonic fibroblasts (MEFs) in hESC medium are likely in a partially primed pluripotent (PPP) state with a heterogeneous expression of naïve-specific markers.

**Figure 5.**
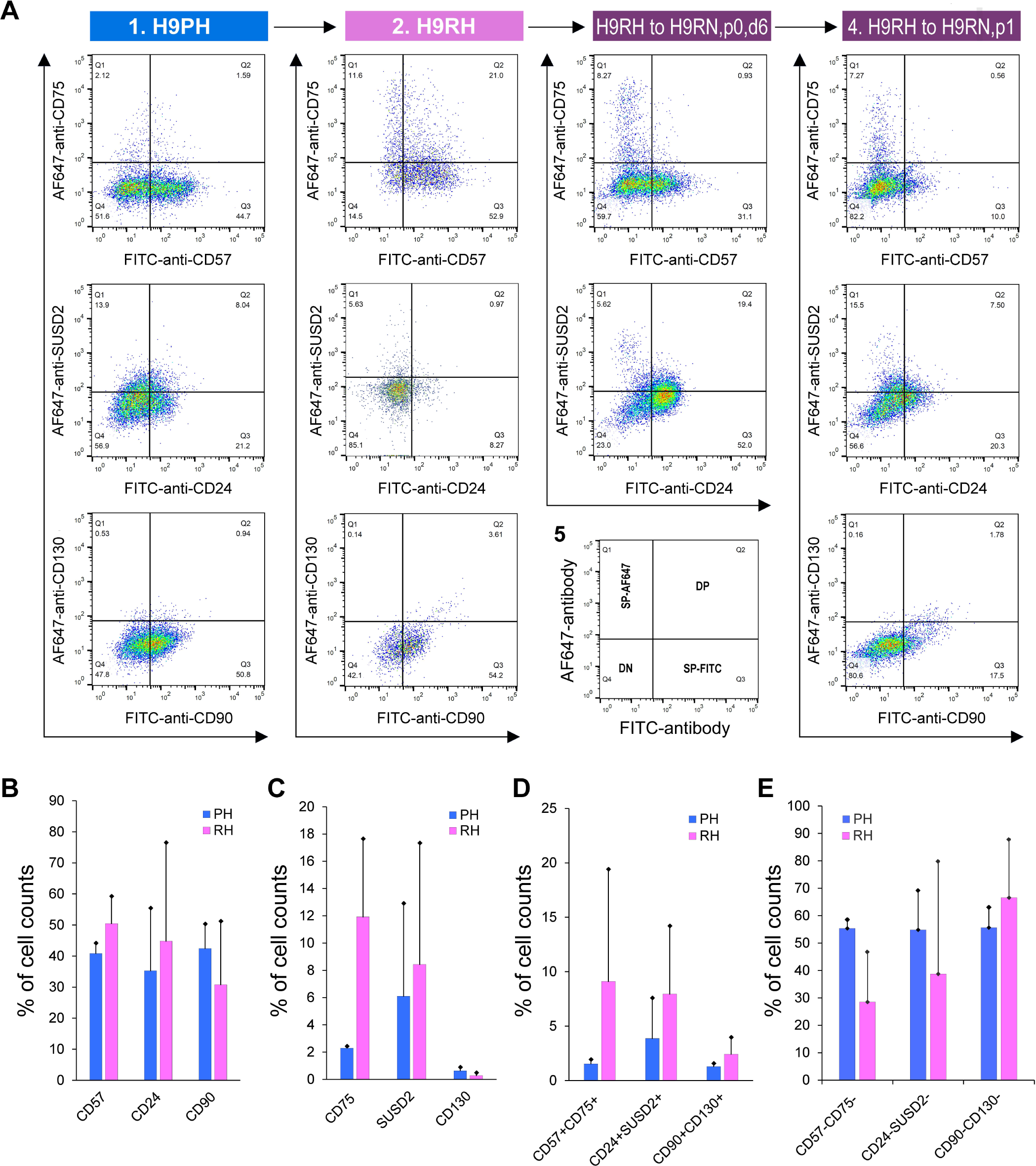
Flow cytometric analysis of primed- and naïve-specific surface expression. **(A)** Expression of the primed-specific markers (CD57, CD24, and CD90) as well as the naïve-specific surface markers (CD75, SUSD2, and CD130) in Hoechst 33258-gated primed and RSeT H9 cells under hypoxia (H9PH and H9RH respectively), H9RH cells grown under normoxia for 6 days (i.e., H9RN, p0, d6) and 1 passage (H9RN, p1). (**B to D**) Statistical analysis of the primed-specific markers (CD57, CD24, and CD90), naïve-specific surface markers (CD75, SUSD2, and CD130), double positive and double negative subsets in Hoechst 33258-gated H1, H7, and H9 single cells. Data are represented as the mean ± SD (standard deviation) of fluorescence intensity in H1, H7, and H9 lines (n = 3). PH, primed hESCs grown under hypoxia; RH, RSeT-converted hESC grown under hypoxia.

### RSeT hESCs are deficient in the expression of naïve-specific surface markers

To further examine the naïve state in RSeT hESCs, we employed the same set of primed- and naïve-specific markers to assess the heterogeneity of RSeT cells (Figures 4B-D, 5B, 5C). When primed hESCs were switched to RSeT medium, we observed a 15% increase in CD75, 17% in SUSD2, and 0.1% in CD130 expression in RSeT H1 cells (Figures 4B-D, second row). Evidently, only a 4% increase was found in CD75, with no increase in both SUSD1 and CD130 expression in RSeT H7 cells (Figure 4B-D, the fourth row). In the case of H9 cells, there is a 9.5% increase in CD75 in RSeT H9 cells, concomitantly with decreased SUSD2 (Figure 5A, middle panel). Moreover, the expression of CD75 tended to decrease under the normoxic conditions (Figure 5A, top panel).

To gain a deeper understanding of the transitions from primed to naïve states, we examined the changes in three pairs of double positive subpopulations, CD57^+^CD75^+^, CD24^+^SUSD2^+^, and CD90^+^CD130^+^. These three pairs were elevated by 7.6%, 4.1%, and 1.1%, respectively, in RSeT hESCs when compared with primed controls (n = 3 cell lines) (Figure 5D). Taken together, our data suggest that only a subset of H1 cells responded to naïve conversion with RSeT medium. However, the CD75^+^ and/or SUSD2^+^ subsets in RSeT H1 cells failed to express the LIF coreceptor CD130/IL6ST (Figure 4D). Hence, the RSeT medium has a limited ability to reprogram primed hESCs into a complete naïve state (Figures 5B-E). This might be due to the absence of some core signaling transduction pathways (e.g., the LIF coreceptor pathway) required to sustain the naïve state.

### Primed hESCs display differential sensitivity to the inhibition of core signal transduction pathways

To determine the response of both primed and RSeT hESCs to core signaling factors, we initially evaluated primed hPSC growth patterns in three hESC lines (H1, H7, and H9) under both normoxic and hypoxic conditions. We found that hypoxia did not significantly increase hESC colony numbers (data not shown), suggesting that hypoxia alone is not a major factor that affects the growth patterns of primed hESCs. It is well known that primed hPSCs require FGF2/TGFβ/Nodal/Activin A for cell proliferation and self-renewal (James et al., 2005; Thomson et al., 1998; Vallier et al., 2005; Xu et al., 2005). It is expected that some core signaling pathway-related growth factors might regulate hPSC growth potential.

We next examined the requirements of some key signaling pathways (e.g., FGF2 and TGFβ) in primed hESCs under both normoxic and hypoxic conditions (Figures 6A-C). Inhibition of FGF2 by its specific inhibitor, PD173074, indeed resulted in differential colony growth in the three hESC lines, in which H1 cells were sensitive to FGF2 inhibition under both normoxic and hypoxic conditions (Figure 6A, lanes 1-4, 13-16). H7 cells showed a moderate FGF2 inhibition under the above two conditions. However, H9 cells were susceptible to FGF2 inhibition only under <u>P</u>rimed and <u>N</u>ormoxic growth conditions (PN) (Figure 6A, lanes 9-12). This inhibitory pattern in H9 cells was partially restored by the FGF2 inhibitor under the <u>P</u>rimed and <u>H</u>ypoxic growth condition (PH) (Figure 6A, lanes 21-24).

**Figure 6.**
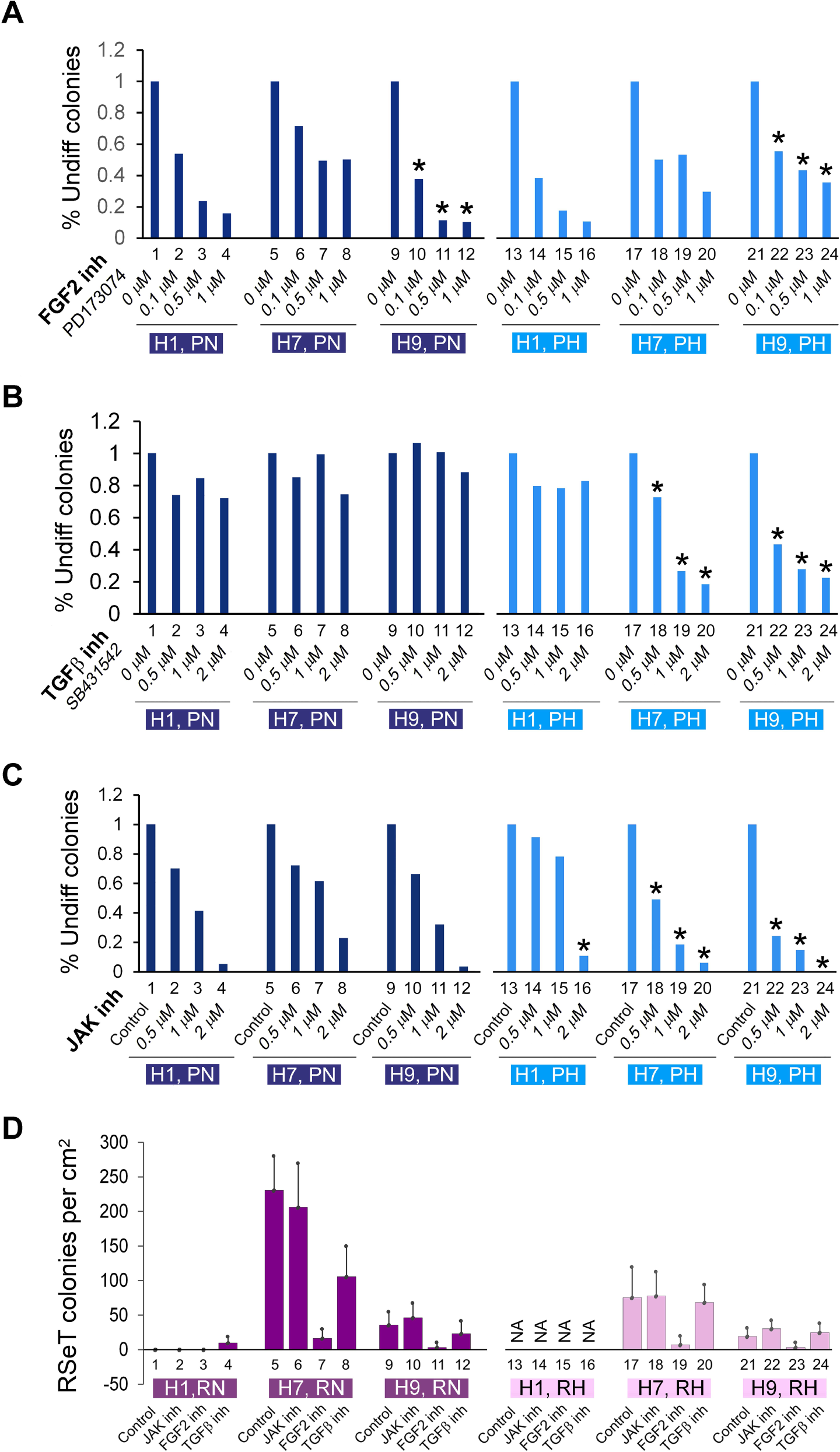
Single-cell colony growth assays of FGF2-TGFβ-, and JAK-dependent hESC colonies. (**A**-**C**) Primed H1, H7, and H9 cells, grown on MEFs under normoxic and hypoxic conditions, treated with the FGF2 inhibitor PD173074, TGFβ1 inhibitor SB431542, and JAK inhibitor I (JAKi) as indicated. The hESC colonies were fixed at day 3 in 4% formaldehyde, stained for alkaline phosphatase positive (undifferentiated) colonies, and counted. The asterisk signs indicate the major changes observed between primed and/or RSeT cell lines. (**D**) JAK, TGFβ, and FGFR pathway inhibition assays in RSeT hESC lines, with JAK inhibitor I (0.5 μM), the FGF2 inhibitor PD173074 (0.1 μM), and the TGFβ inhibitor SB431542 (0.5 μM). The cell colonies were stained with the anti-NANOG antibody. Data are represented as the mean ± SD (standard deviation) of NANOG positive colonies counted randomly at 5 regions (n = 5) under an immunofluorescence microscope. inh, inhibitor or inhibition; NA, cells not available at the time of the assay due to the significantly slow growth of cells.

With respect to the effect of TGFβ inhibition, the three hESC lines exhibited similar growth patterns under PN conditions, and relatively insensitive to TGFβ inhibition mediated by the TGFβ inhibitor SB431542, in the range of 0.5-2 μM range (Figure 6B, lanes 1-12). Interestingly, H1 cells appeared not to be affected by TGF-β inhibition in both PN and PH conditions, an unexpected result that has not been previously reported (Figure 6B, lanes 1-4, 13-16). However, both H7 and H9 cells showed a dose-dependent TGFβ inhibition only under PH (Figure 6B, lanes 17-24).

To examine the intrinsic naïve growth capacity in primed hESCs, we performed a Janus kinase I (JAK) inhibitory experiment under the assumption that naïve ES colonies (e.g., mESCs) would be greatly reduced or eliminated due to their hypersensitivity to JAK inhibition (Tesar et al., 2007). As shown in Figure 6C, the three hESC lines showed a similar JAK inhibition under 0.5 μM JAK inhibitor I (JAKi) treatment (Figure 6C, lanes 1-12). However, H1 exhibited much lower sensitivity to JAKi under PH, which is in contrast with both H7 and H9 cells, which manifested an enhanced response to JAKi (Figure 6C, lanes 13-24).

Collectively, we identified several different intrinsic properties peculiar to these three primed hESC lines. The differential requirement of FGF2 in primed hESCs is both cell-line- and oxygen-tension-specific. Compared with both H7 and H9, H1 cells are relatively sensitive to FGF2 inhibition and resistant to JAKi, two of the hallmarks of primed pluripotency. These data suggest that the intrinsic naïve-compatible activity is higher in both H7 and H9 than H1 cells. This differential naïve signaling compatibility might enable H7 and H9 cells to readily adapt themselves in some naïve-like growth media such as RSeT medium (Figure 1C, middle and bottom panels). Thus, the intractability of primed H1 cells to be induced and maintained as domed colonies might also be consistent with the lower intrinsic JAK activity, which is associated with the LIF-CD130/IL6ST-JAK pathway (Collier et al., 2017) in these H1 cells (Figure1C, top panel; Figure 6C, lanes 13-15).

### Heterogeneous growth of RSeT hESCs depicted by their differential sensitivity to the inhibition of core signal transduction pathways

In parallel, we also determined the response of RSeT H1 cells to core signaling factors in SCPE assays. Consistently, H1 SCPE under RN conditions was undetectable (Figure 6D, lanes 1-3). It was also difficult to obtain H1 cell colonies under RH conditions for a comparative analysis in this assay due to their extremely slow growth rate (Figure 6D, lanes 13-16). Noticeably, under normoxia, both H7 and H9 RSeT cells were completely resistant to JAK inhibition, highly sensitive to FGF2 inhibition, and moderately sensitive to TGFβ inhibition (Figure 6D, lanes 5-12). Similar signaling response patterns were also observed in these RSeT cells under hypoxia (i.e., RH conditions), except that the partial sensitivity to TGFβ inhibition in these cells was abrogated, comparable with the levels of untreated controls (Figure 6D, lanes 17-24). Of note, compared with hypoxia, a significantly higher number of colonies formed in RSeT H7 cells under normoxia (Figure 6D: lanes 5-8, 17-20), further demonstrating the hypoxia-independent cell growth feature in RSeT hESCs.

## DISCUSSION

Increasing evidence suggests the existence of a broad spectrum of pluripotency between naïve and primed pluripotency among different mammalian species (Rossant and Tam, 2017; Wu et al., 2017). Various hPSC models derived *in vitro* cell culture systems may echo these distinct pluripotent states (Figures 1A, 3, and 7). It is possible to achieve a naïve pluripotent state in hPSCs, as indicated in both t2iLGo and 5iLA models (Figures 2C, 7A). The t2iLGo protocol utilizes titrated 2iL in the presence of the PKC inhibitor Go6983 and transgene expression of *KLF2* or *NANOG* (Takashima et al., 2014) or OSKM (i.e., the combination of OCT4, SOX2, KLF4, and c-MYC) (Liu et al., 2017) (Figure 3C). In a revised version of t2iLGo without transgene expression, known as PXGL (Bredenkamp et al., 2019; Guo et al., 2017), GSK3 inhibition is replaced by the tankyrase inhibitor, XAV939. This modified version is now being widely used to promote the establishment of the human naïve pluripotent state (Kagawa et al., 2022; Khan et al., 2021; Zhou et al., 2023). Moreover, the 5iLA protocol evolved from the 2iL protocol into a five-inhibitor-based protocol, which impedes MEK, GSK3, ROCK, BRAF, and SRC pathways (in the presence of LIF and activin), to orchestrate profound genomic and epigenomic changes in primed hPSCs (Collier et al., 2017; Theunissen et al., 2016).

**Figure 7.**
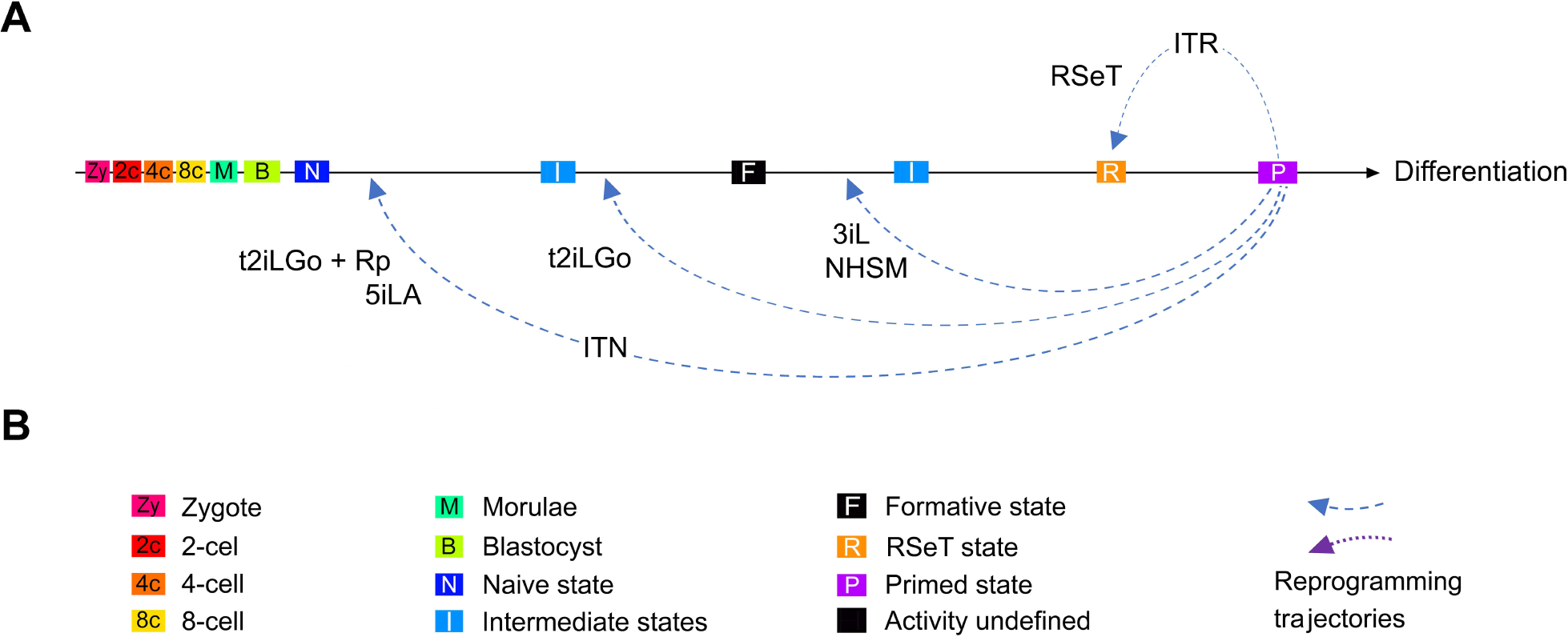
Regulation of pluripotent states by integrated signal inputs from growth media. **(A)** Reprogramming the primed pluripotent stem cells to naïve, formative, and RSeT-based states. Differential growth signaling molecules in media may limit the reprogramming efficiency and trajectories observed in different naïve conversion protocols such as t2iLGo plus reprogramming transgenes (t2iLGo + Rp), 5iLA, t2iLGo, 3iL, NHSM, and RSeT, resulting in diverse pluripotent states upstream or downstream of the formative state. (**B**) Color key, symbols, and abbreviations for cell types, cellular or pluripotent states, and integrated reprogramming trajectories of <u>n</u>aïve (ITN) or <u>R</u>SeT (ITR) states.

However, several naïve growth methods only enable the emergence of some LIF-dependent signatures similar to those of native preimplantation epiblasts (Chan et al., 2013; Cornacchia et al., 2019; Gafni et al., 2013; Valamehr et al., 2014; Zimmerlin et al., 2016). Of note, without the expression of powerful transgenes, the naive conversion with t2iLGo would be compromised, possibly ending up with a primed-to-naïve intermediate state (Figure 3C) (Guo et al., 2016). The pluripotent states in these hPSC models are fundamentally similar to the executive phase of pluripotency, coined as the formative state (Smith, 2017).

In this study, we have redefined and characterized RSeT-based hESCs, which exhibit several new features that have not been previously characterized. These features include hypoxia-independent cell growth, a unique pluripotent state with heterogeneous expression of primed- and naïve-specific markers, differential dependency of FGF2, JAK, and TGFβ signaling in a cell-line-specific manner. These prominent characteristics distinguish these cells from those of primed and naïve hPSCs, which warrants further discussion as follows.

The converted hESCs appear to acquire a specific state downstream of the formative state with differential growth potentials in individual RSeT hESC lines (Figures 3C, 7A). Interestingly, either RSeT medium-converted (cRSeT) cells or RSeT plus transgene (OKSM) reprogrammed (rRSeT) cells show global similarity to primed hESCs (Figure 3C) (Johnson et al., 2021; Liu et al., 2017). Also interestingly, rRSeT cells are resistant to transgenic reprogramming effects, suggesting that the restricted reprogramming capacity in RSeT hPSCs is difficult to circumvent even in the presence of powerful transgene expression. These data further support the unique pluripotent state in RSeT cells, which possess limited pluripotent plasticity and lower naive reprogramming efficiency.

Thus, the unique pluripotent state in RSeT hESCs might help to address several fundamental developmental questions on the divisibility of the pluripotent window and the state plasticity of hPSCs. Indeed, RSeT hESCs represent a novel state characterized by FGF2-dependency and JAK- and TGFβ co-independency (Figure 6), between the formative and primed states, as evidenced by our detailed characterization and genome-wide profiling (Figures 1-6). It was also shown that RSeT cells did not undergo X chromosome reactivation, which was unveiled by RNA FISH, a more conclusive assay (Vallot et al., 2017). Consistently, we reveal that RSeT hESCs do not acquire many naïve pluripotency hallmarks such as global hypomethylation and transcriptomic signatures as documented in multiple naive protocols.

Additionally, substantial differences between laboratories in the way they handle primed hPSC cultures can significantly affect the outcomes of converting them to naïve states in vitro and make data interpretation confusing (Johnson et al., 2019). The H9 cell line is an example of this (Johnson et al., 2021). H9 cells have been widely used for hPSC genetic engineering and lineage differentiation (www.wicell.org) (Johnson et al., 2021; Mallon et al., 2013; Takashima et al., 2014; Wang et al., 2011). Primed H9 cells might have acquired substantial genetic and epigenetic changes to adapt themselves to various pluripotent states or differentiation conditions. Hence. it is conceivable that H9 cells exhibit much higher growth advantage and adaptability under various RSeT conditions. A high-resolution genome sequencing may provide a definite answer in the future. The primed hESC lines used in this study have a PPP state and closely resemble primary hESCs. This might reflect a more physiological state of hESCs in vitro. Therefore, hESCs with the PPP state are especially suitable for hPSC research.

In conclusion, we have redefined RSeT as FGF2-dependent and JAK- and TGFβ co-independent cellular models, with a unique pluripotent state between formative and primed pluripotency. RSeT cells are relatively insensitive to hypoxic tension. Primed hPSCs may have a restricted plasticity to be converted to the formative and naïve states under certain hPSC growth conditions (such as RSeT). It is imperative to have a thorough understanding and mastery of physiologically relevant growth conditions in order to maintain desired pluripotent states for disease modeling, drug discovery, and future clinical applications.

## Materials and Methods

### Cell lines

Three NIH registered human embryonic stem cell (hESC) lines, H1 (WA01, registration number 0043), H7 (WA07, registration number 0061), and H9 (WA09, registration number 0062), were used in this study. All three hESC lines were obtained from WiCell Research Institute (Madison, WI) and previously extensively characterized in our NIH Stem Cell Unit (Chen et al., 2014; Chen et al., 2012; Mallon et al., 2013). More detailed information regarding these hESC lines is available (stemcells.nih.gov/research/).

### MEF isolation and cell culture

MEFs were isolated from the mouse CF1 strain and grown on 0.1% gelatin-coated culture plates in DMEM/F12 medium supplemented with 10% fetal bovine serum (Gemini Bio-Products, West Sacramento, CA), 2 mM L-glutamine, and 0.1 mM non-essential amino acids.

### hESC culture on MEF feeder

All hESC lines were initially cultured according to the supplier’s protocols and adapted to a simple protocol as outlined below. The MULTIWELL^TM^-6 WELL polystyrene plates were coated with 0.1% gelatin for 1 hour at 37°C. MEFs, at passage number 5 and 6 (designated as p5 and p6, respectively), were irradiated at a dose of 8,110 rads with an X-ray machine (Faxitron Bioptics, LLC., Tucson, AZ). The irradiated cells were plated at a density of 2 x 10^4^ cells/cm^2^ and incubated at 37°C for 24 hours. HESCs were plated on MEF feeder layer as small clumps (∼ 50 to 100 µm in diameter) in the hESC medium containing 80% DMEM/F12 medium, 20% knock-out serum Replacement (KSR), 2 mM L-glutamine, and 0.1 mM non-essential amino acids, 0.1 mM β-mercaptoethanol, and 4 ng/mL of FGF2. Detailed descriptions about hESC passage, expansion, and cryopreservation are available in our previous reports (Chen et al., 2014; Chen et al., 2012; Mallon et al., 2013).

### RSeT conversion, passages, and maintenance

RSeT medium has been advertised as a “defined” medium (StemCell Technologies Inc. Vancouver, catalog number 05970), without the inclusion of both FGF2 and TGFβ. RSeT converts and maintains primed hPSCs in a “naïve-like” state (www.stemcell.com). This medium was based on NHSM (Gafni et al., 2013), but made by a different formulation under a license from the Weizmann Institute of Science (Israel). The primed H1, H7, and H9 hESCs were converted to a new pluripotent state on MEFs in RSeT medium as recommended by the vendor’s protocol as described below.

After 3-day induction or selection of primed hESCs, the cell colonies (in each well) were dissociated as single-cell suspension by TrypLE™ Express (Thermo Fisher Scientific, REF number 12604-013) and passaged on MEF feeder in 6-well plates in a 1:6 split ratio. The cells were grown in a humidified incubator with 5% CO_2_, 3% O_2_ (hypoxia), or 20% O_2_ (normoxia). The ROCK inhibitor Y-27632 (10 µM) was added to enhance SCPE for the initial 3 passages. The RSeT cell lines were fully characterized at passage 10 in this study.

Primed and RSeT cell growth, proliferation, and pluripotency assays were examined in downstream experiments (or analyses) such as alkaline phosphatase, immunofluorescence assays, microarray, real-time PCR, and cell growth assays. RSeT cell lines were routinely analyzed with their primed counterparts. We refer to samples from different cell lines (e.g., H1, H7, and H9) with different passages (batches) as biological replicates and those triplicate or quadruplicate determinants in one independent experiment as technical replicates in this study.

### Clonogenicity of primed and RSeT hESCs under the inhibition of JAK, TGFβ, and FGFR pathways

RSeT lines (at passage number 15) were dissociated in TrypLE solution, counted, and seeded at a density of 3 x 10^4^ cells per well (in 12-well plate) on MEF feeder in 0.5 mL of RSeT complete medium. Drugs and growth factor-containing medium were added next day, incubated for 96 hours, and then cells in 4% formaldehyde. The cell colonies were stained with anti-NANOG antibody and 5 regions were randomly counted at in three wells (technical replicates) under an immunofluorescence microscope. Data are represented as the mean ± SD (standard deviation).

### Alkaline phosphatase assay

Alkaline phosphatase staining (Alkaline Phosphatase Staining Kit II, catalog number. 00-0055, Stemgent, CA) was used to assess the number of pluripotent stem cells and their differentiated or partially differentiated counterparts. Cells were fixed in 4% formaldehyde for 20 minutes and permeabilized in 0.05% Tween-20 in D-PBS for 5 minutes at room temperature. Cells were then incubated with the AP Staining Solution A, B, and mix at room temperature for 20 minutes followed by rinsing cells twice in D-PBS. All samples were analyzed in the presence of D-PBS. Red, purple, and dark black colored colonies, which were NANOG and OCT4 positive, were counted as pluripotent colonies. Data are represented as the mean ± SD (standard deviation).

### RNA preparations

RNA was extracted by using the basic Trizol protocol (Invitrogen Inc.) with slight modifications. Briefly, hESCs per sample were dissolved in 1 mL of the Trizol solution and stored in −80°C freezer prior to use. HESC lysates were further subjected to chloroform extraction and isopropanol precipitations. A DNA-Free kit (Applied Biosystems/Ambion, Austin, TX) was used to remove genomic DNA contamination. The quality of RNAs was agarose gel-verified and the concentration determined by using the ND-1000 UV-Vis Spectrophotometer (NanoDrop Technologies Wilmington, DE).

### Complementary DNA (cDNA) microarray

Genome-wide gene expression analysis in primed and RSeT hESCs (at passage 10) was performed and analyzed with the Agilent chip (Agilent Technologies Inc., Santa Clara, CA). Global gene expression microarray was performed and analyzed using Agilent software, reagents according to the manufacturer’s instructions (Mallon et al., 2013). Sample amplification and labeling were carried out using the Low Input Quick Amp labeling Kit (Catalog number 5190-2305, Agilent Technologies, Santa Clara, CA). Briefly, total RNAs (200 ng) were reverse transcribed into cDNAs using T7 primer mix in 5X First Strand Buffer, 0.1 M DTT, 10 mM dNTP Mix, and Affinity Script RNase Block Mix. The cDNAs were used to synthesize Cy3-labeled cRNAs in a reaction containing nuclease-free water, 5X Transcription Buffer, 0.1 M DTT, NTP Mix, T7 RNA polymerase blend, and Cyanine 3-CTP. The RNeasy Mini Kit (QIAGEN) was used to purify the linearly amplified cRNA samples. Cy3-labeled cRNAs were measured using the NanoDrop Spectrophotometer.

Cy3-labeled cRNAs were further hybridized onto Agilent Whole Human Genome kit 4 x 44K slides (Catalog number G4112F, Agilent Technologies) containing 44,397 oligonucleotide probes in 2X Hi-RPM Hybridization buffer (Catalog number 5188-5242, Agilent Technologies) at 65°C for 17 hours at 10 r.p.m. as recommended. Slides were scanned using an Agilent DNA microarray compatible scanner, with a one-color scan setting for 4 x 44k array slides. Quantification files were processed using Agilent Feature Extraction Software. We also used quantile normalization and background correction for further data analysis.

### Quantitative real-time PCR (qPCR)

For qRT-PCR, genomic DNA-free RNAs (2 μg) were reversely transcribed (in a 20-μl volume) into cDNAs using the SuperScript™ VILO™ cDNA Synthesis Kit (Thermo Fisher catalog number 11754) and diluted in 380 μl nuclease-free H2O. All TaqMan™ MGB probes were purchased from Thermo Fisher Scientific Inc. (see Key Resource Table). Both β-actin (*ATCB*) and glyceraldehyde-3-phosphate dehydrogenase (*GAPDH*) assays were used as internal controls for normalization. Approximately, 100 ng of cDNAs were amplified in 1X TaqMan™ Fast Advanced Master Mix and 1X TaqMan™ Assay in a 20-µL reaction using the QuantStudio 6 Flex real-time PCR Platform (Thermo Fisher Scientific Inc.) according to the manufacturer’s instructions. Thermal cycling was done with 2 minutes at 50°C, 20 seconds at 95°C followed by 40 cycles of PCR amplification (i.e., 1 second denature at 95°C and 20 seconds at 62°C). Equal amplification efficiencies in TaqMan assays were confirmed by serial cDNA dilutions. Data were analyzed by a three-step comparative cycle threshold (Ct) method, which includes (1) normalization to endogenous control (Ct_Target gene_ – Ct_Endogenous control_ = ΔCt), (2) normalization to calibrator sample (Ct_Sample_ – Ct_calibrator_ = ΔΔCt), and (3) the final use of formula (2^-ΔΔCt^) to calculate the relative fold change of gene expression.

### Flow cytometry

Flow cytometric analysis of conventional pluripotent stem cell surface markers for primed and RSeT hESCs was initially performed as described previously (Mallon et al., 2013). For primed- or naïve-specific cell surface marker expression, we modified the above method as follows. Briefly, hESC colonies were grown on MEF feeder in hESC and RSeT media in a 6-well plate. For hESCs on feeders, the cells were rinsed once in 1X D-PBS and dissociated by Collagenase type IV for 40 min. Only floating colonies were collected, sedimented once in D-PBS, further digested in 1 mL of TrypLE for 10 min into single-cell suspension followed by single cell confirmation under the microscope. The enzymatic activity was inactivated by adding FACS buffer (DMEM/F12 medium containing 10% FBS). Then, 2 mL of cells (∼1 x 10^6^ cells) were transferred into a Falcon 5 mL-polypropylene round-bottom tube (REF 352063, Corning) and centrifuged in a swing bucket at 300 *g* for 5 minutes. The above wash step with FACS buffer was repeated once. The cell pellets were resuspended in 100 µL of FACS buffer and incubated with Alexa Fluor® 647 (AF647)- or FITC-conjugated primary antibodies or isotype controls (4 μg/mL, Key Resource Table) on ice for 1 hour with gentle flickering the tubes every 20 minutes. The incubation was terminated by adding 4 mL of FACS buffer followed by centrifugation at 300 *g*. The cell pellets were then resuspended in 0.5 mL of D-PBS and stained with 2.5 μL of bis benzamide (Hoechst 33258) solution (0.2 mg/mL, Sigma) at room temperature for 20 minutes. The cells were further fixed (in 4% of formaldehyde solution) for 20 minutes at room temperature. The fixative was removed by rinsing the cells twice in D-PBS followed by filtering the cell solution into a Falcon 5-mL polypropylene round-bottom tube with a cell-strainer cap (REF 352253, Corning). Finally, the cells were analyzed by a Beckman Coulter machine at the NINDS Core Facility (National Institutes of Health, Bethesda, MD) or by a Guava easyCyte™ device (Luminex Corporation, Austin TX). We used DAPI 405-448/59-area (at X-axis) versus Y-axis DAPI-405-448/59-width (at Y-axis) to gate nucleated single cells for data analysis by FlowJo v10.9.0 (www.flowjo.com). All positive staining cells are related to their isotype (negative) controls, which are included in every independent experiment. These controls are partially shown in Figure S4.

### Immunofluorescence

Primed and RSeT hESCs were grown either in 6- or 24-well plates and fixed in 4% formaldehyde at room temperature for 20 minutes. The cells were also grown on ibidi chamber slides (ibidi USA, Inc., Fitchburg, WI) for confocal microscopic analysis. Cell samples were blocked with 10% normal goat serum and 0.1% bovine serum albumin (BSA) in the presence or absence of 0.1% Triton X-100 (diluted in D-PBS) at room temperature for 1 hour, then reacted with primary antibodies at 4°C overnight or at room temperature for 2 hours, and subsequently incubated with Alexa Fluro® conjugated secondary antibodies (Thermo Fisher Scientific Inc., Waltham, MA) in 5% blocking solutions for 1 hour at room temperature. Finally, the cells were stained with Hoechst 33258 solution or DAPI. All images used for quantitative analysis were obtained under unsaturated exposure conditions. The detailed procedures are also available in our previous reports (Chen et al., 2014; Chen et al., 2012; Mallon et al., 2013).

### Microscopy

Cell samples were examined under a series of Zeiss microscopes, including Axiovert 25 light microscope, Axiovert 200 fluorescence, Axio Observer, and LSM 810 confocal microscopes (Zeiss, Jena, Germany). These microscopes were equipped with corresponding software such as Adobe Photoshop® (Adobe Inc., Mountain View, CA), AxioVision Rel 4.6 (Zeiss), and ZenBlue (Zeiss) for image acquisitions and analysis.

### Microarray data analysis

The microarray data analysis was performed essentially as described previously (Chen et al., 2012; Mallon et al., 2013). ANOVA (analysis of variance) was used to assess the number of genes (probes) (n = 43,376) enumerated in the datasets, in which 32,563 probes had the expression values greater than background, 26,317 probes had the values greater than the noise cut-off value 6.5 in at least one sample, and 1,958 probes showed type III ANOVA corrected *P*-values less than 0.05. Gene probes selected for use represent those without noise-, processing-, and gender-biased ones. These probes passed multiple tests, including ANOVA under multiple comparison correction condition (*P* < 0.05) using class as the factor, *post-hoc* tests for at least one pair-wise class comparison (Tukey HSD *P* < 0.05), and a mean/difference criterion for the same pair-wise class comparison having a Tukey HSD *P* < 0.05. Pearson correlation heatmap depicts 12 cell lines using log (base = 2) transformed and quantile normalized mRNA expression. PCA was performed as described previously (Mallon et al., 2013).

### Meta-analysis of human pluripotent stem cell (hPSC) and primate early embryo datasets

Comparative meta-analysis of hPSC datasets generated from various protocols was performed by integrating three interrelated analytic tools including PCA, *t*-SNE, and SC3 consensus clustering methods, which were described previously (Johnson et al., 2021). This multivariate meta-analysis platform consists of a series of independent modules that constitute a gene expression matrix, which comprises 16 datasets, including the microarray dataset D21 in this study.

All naïve protocols were based on the modifications of the 2iL protocol [i.e., MEK and GSK3 inhibitors in the presence of leukemia inhibitory factor (LIF)]. The related datasets (with GSE or EMBL-EBI accession numbers in parentheses) include: D3 (E-MTAB-2031) from 3iL: MEK, GSK3, and BMP pathway inhibitors in the presence of LIF in TeSR1 medium (Chan et al., 2013); D5 (GSE46872) from NHSM (Gafni et al., 2013); D6 (GSE50868) fate maintenance medium (FMM) (Valamehr et al., 2014); D7 (GSE59435) from MEK, GSK3, ROCK, BRAF, and SRC pathway inhibitors in the presence of LIF and Activin (5iLA) (Theunissen et al., 2016; Theunissen et al., 2014); D22B (E-MTAB-2857) from titrated MEK and GSK3 inhibitors with LIF plus the PKC inhibitor Go6983 (t2iLGo) (Takashima et al., 2014); D23 (E-MTAB-2856) from t2iLGo (Takashima et al., 2014); D24 (E-MTAB-4461) also from t2iLGo (Guo et al., 2016); D25 (GSE36552) containing human embryo datasets (Yan et al., 2013); D26 (GSE29397) representing the second human embryo dataset (Vassena et al., 2011); D27 (SRP115256) encompassing a panel of datasets from multiple naïve protocols at the same laboratory (Liu et al., 2017); D28 (GSE44430) from 2iL in the presence of the tankyrase inhibitor XAV939 (XAV939/2iL) (Zimmerlin et al., 2016); D29 (GSE141639) from XAV939/2iL (Park et al., 2020); D30 (GSE66507) containing human blastocysts at embryonic day 6 to 7 (E6-E7) (Blakeley et al., 2015); D31 (E-MTAB-3929) having human embryos at E3-E7 (Petropoulos et al., 2016); and D32 (GSE74767) encompassing Cynomolgus monkey embryos at E7-E17 (Nakamura et al., 2016).

To integrate the *<u>Cynomolgus</u>* monkey dataset D32, Fastq files for 34 *<u>Macaca fascicularis</u>* samples were downloaded from NCBI SRA (PRJNA301445) and used as input into the nf-core RNA-Seq pipeline (https://nf-co.re/rnaseq/3.10.1, --clip_r1 5 --clip_r2 5) to generate salmon-enumerated gene expression in transcript per million (tpm). The reference genome and annotations used were those for Macaca_fascicularis_6.0 (GCA_011100615.1). Expression was then imported into R (https://cran.r-project.org/), pedestalled by 2, and log2 transformed. To map homologous gene symbols from *<u>Macaca fascicularis</u>* to *<u>Homo sapiens</u>*, biomaRt and SignacX functions were used as described in the “Crabeating_vignette” (https://cran.r-project.org/web/packages/SignacX/vignettes/). All datasets were quantile-normalized, limited to unique genes, percentile-scaled, and batch-corrected (Johnson et al., 2021).

For PCA in meta-analysis, normalized datasets were used to construct a Pearson correlation-based matrix and employed to calculate the principal components using the R programming software. For visualization, PC values were used to map the data points in PCA scatter plots, in which each data point contains the genome-wide gene expression profile of one sample (or cell line). The *t*-SNE analysis was also implemented by the R program using the “Rtsne” function under default parameters (dims = 2, perplexity = 30, theta = 0.5, pca = T, momentum = 0.5, final momentum = 0.8, eta = 200) (Johnson et al., 2021).

## Supporting information

Key Resource Table

## Data and code availability

Data reported in this study will be available in supplemental figures and tables. The microarray data were deposited in the Gene Expression Omnibus (GEO) database at the National Center for Biotechnology Information (NCBI accession: GSE217275). We provide the link below (with a security token: klclsoqmdtuhxuf), www.ncbi.nlm.nih.gov/geo/query/acc.cgi?acc=GSE217275, to facilitate the access to microarray data. Any additional information required to reanalyze the data reported in this study would be available upon request.

## ACKNOWLEDGMENTS

This work was supported by the Intramural Research Program of the National Institutes of Health (NIH) at the National Institute of Neurological Disorders and Stroke (K.G.C., K.R.J., K.P., D.M., Y.C.F., B.S.M.) and the National Institute of Dental and Craniofacial Research (P.G.R).

## AUTHOR CONTRIBUTIONS

K.G.C., B.S.M., and P.G.R.: Conceptualization; K.G.C. and B.S.M.: methodology; K.G.C., B.S.M. P.G.R.: validation and formal analysis; K.G.C., K.R.J., K.P., D.M., F.Y., W.F.L., Y.C.F., B.S.M.: experimentation and investigation; K.G.C.: writing – original draft; K.G.C., B.S.M., P.G.R.; writing – review and editing; K.G.C., K.R.J.: visualization. All authors approved the final manuscript.

