## Supplemental Figures for "Resistance to Naïve and Formative Pluripotency Conversion in RSeT Human Embryonic Stem Cells"

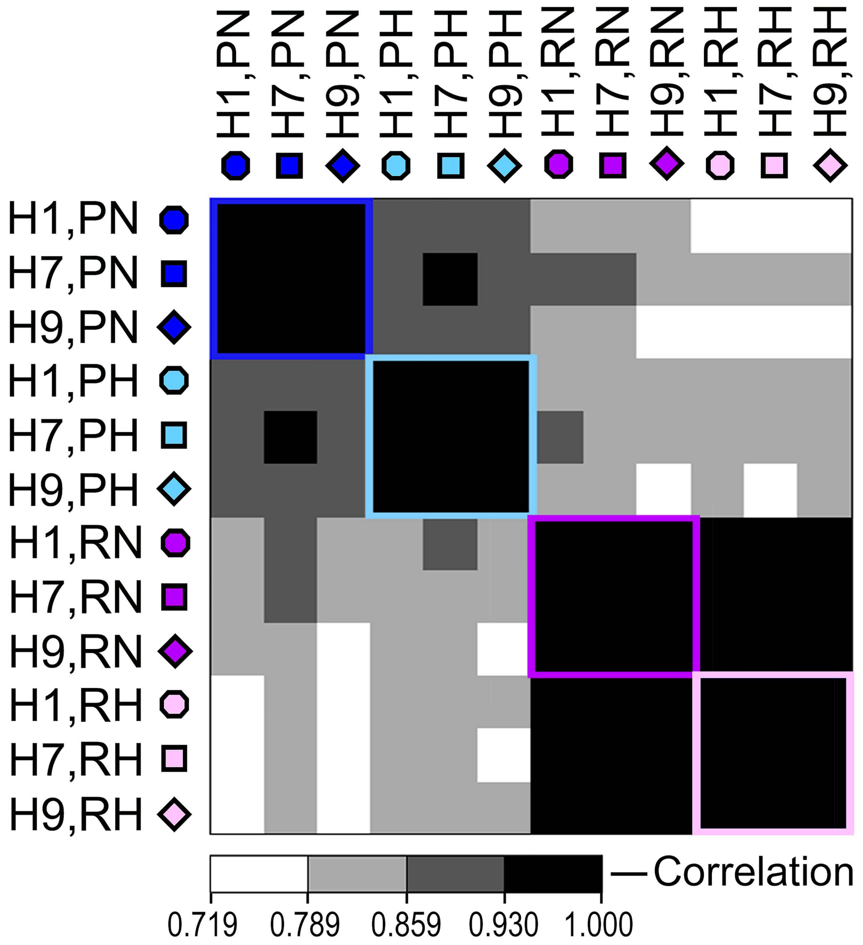


**Figure S1. Pearson correlation heatmap depicts the 12 cell lines using log (base = 2) transformed and quantile normalized mRNA expression (related to Figure 2A).**

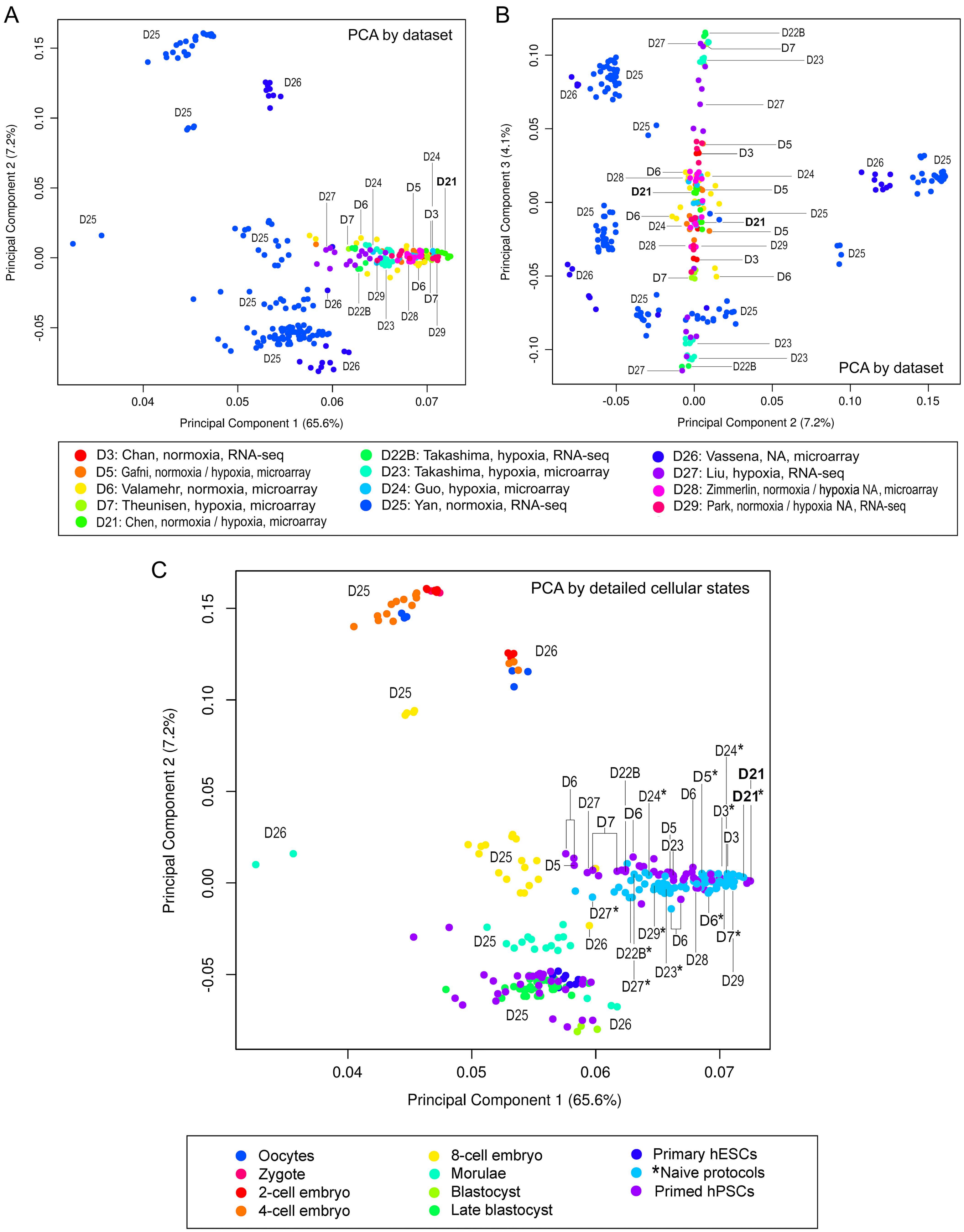


Figure S2. Principal component analysis (PCA) based on datasets and detailed cellular states (Related to Figure 3).

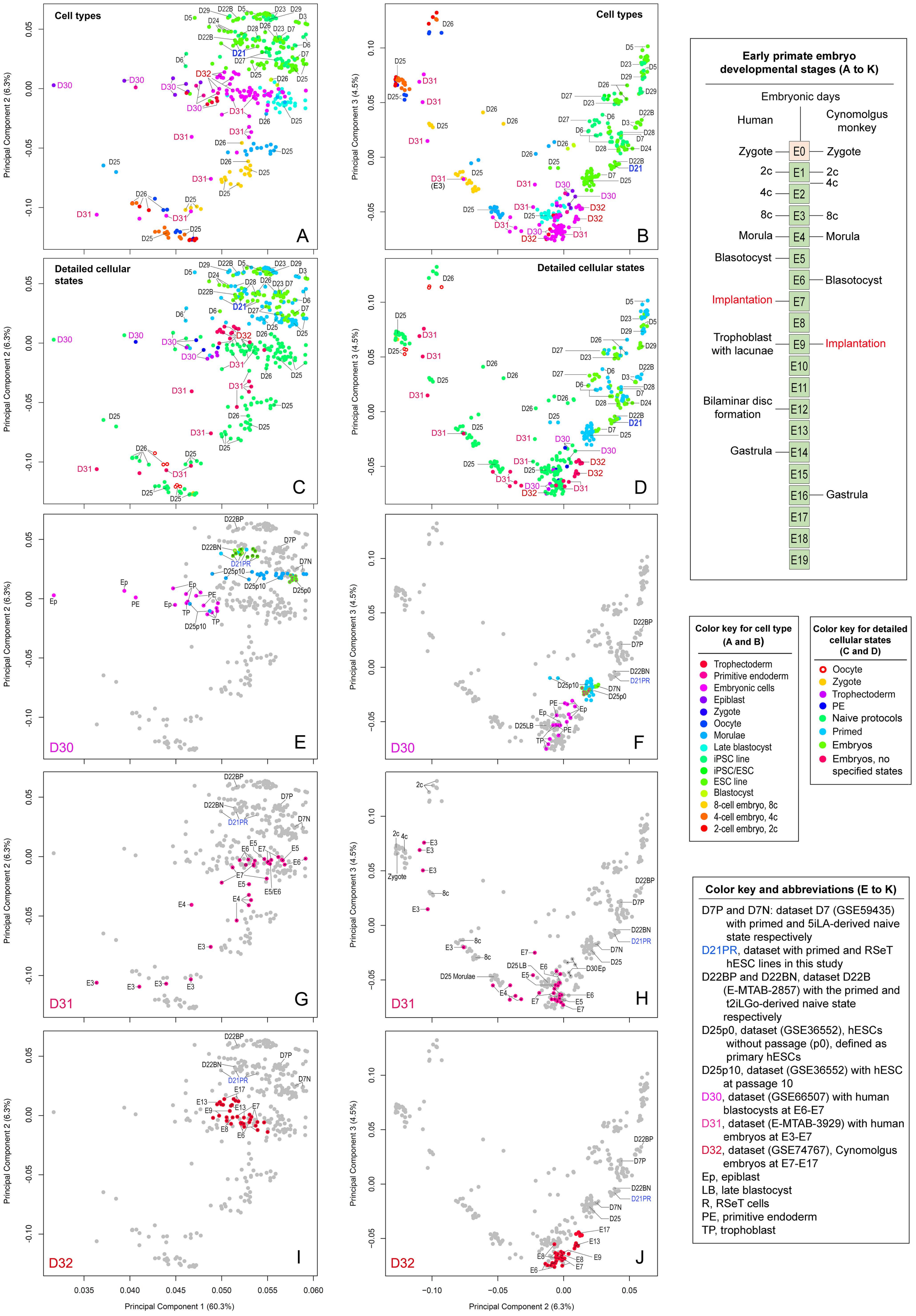


**Figure S3. Principal component analysis (PCA) based on cell types, detailed cellular states, and early embryonic developmental stages (Related to Figure 3A and 3B).**

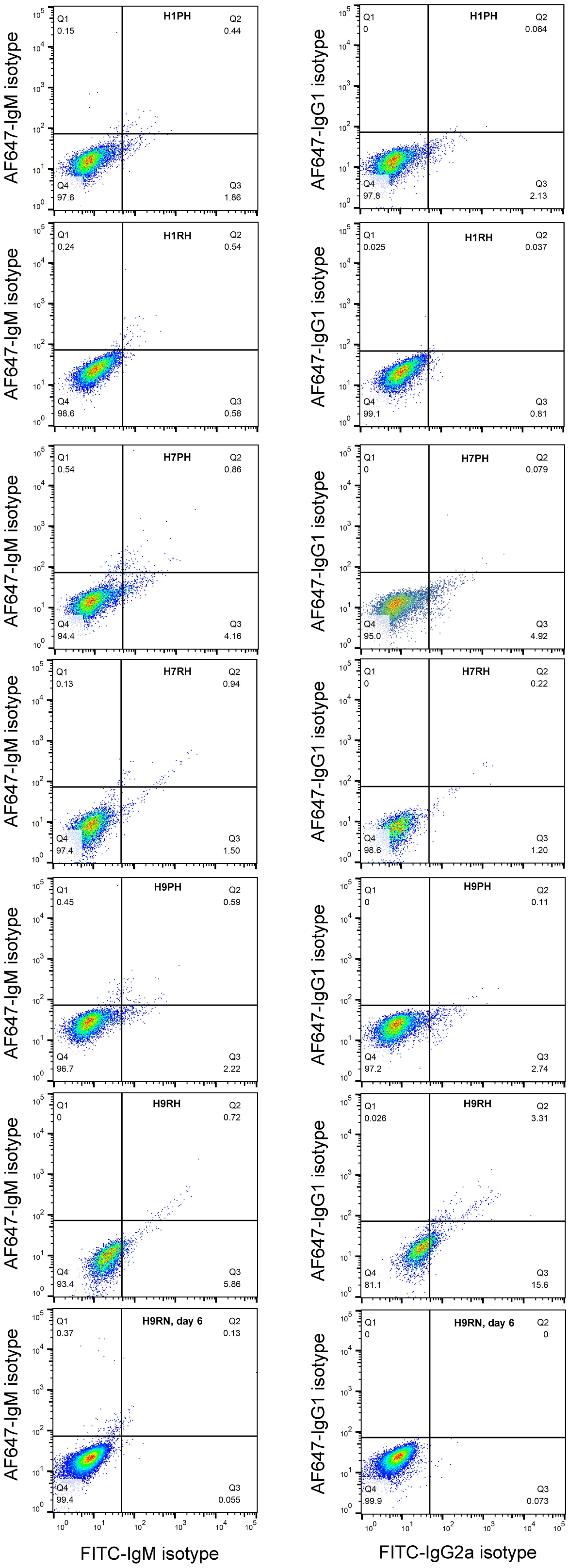


**Figure S4. Representative flow cytometric controls depicting fluorescence labeled isotypoe antibodies used to gate positive antibody staining in primed and RSeT human embryonic stem cell lines (related to Figures 4 and 5)**
