## Supplemental Table 1 for "Resistance to Naïve and Formative Pluripotency Conversion in RSeT Human Embryonic Stem Cells"

### Table S1. Gene expression signatures (n = 112) underlying various pluripotent states

| Gene Symbols | Description (Based on GeneCards and Entrez of NCBI) | Gene expression probes | RSeT vs primed (n = 6, Fold changes) | RN vs PN (n = 3, fold changes) | RH vs PH (n = 3, fold changes) | Notes/ Comments with references[a] |
|---|---|---|---|---|---|---|
| **Inner cell mass (ICM) and other developmental markers not in implantation epiblast** (n = 5, *indicates $P < 0.05$) | | | | | | |
| *ATG2A* | ICM expressed, not in the pre-implantation epiblast, | A_23_P361820 | 1.04 | 0.83 | 1.30 | Pera & Rossant |
| *ATG2B* | ICM expressed, not in the pre-implantation epiblast, | A_23_P88163 | 1.70 | 2.06 | 1.41 | Pera & Rossant |
| *GATA3* | GATA Binding Protein 3, ICM expressed, trophoblast marker | A_23_P75056 | 0.99 | 0.98 | 1.00 | P list in Valamehr, Pera & Rossant |
| *GATA6* | A primitive endoderm marker | A_23_P304450 | 0.19* | 0.08* | 0.42 | Pera & Rossant |
| *MAGEA4* | ICM expressed, not in the pre-implantation epiblast, | A_24_P185945 | 0.96 | 1.00 | 0.92 | Pera & Rossant |
| **Naive-related or epiblast specific gene expression** (n = 60, *indicates $P < 0.05$) | | | | | | |
| *AHNAK* | AHNAK Nucleoprotein | A_24_P943393 | 0.92 | 0.34 | 2.44 | Chan, Epiblast specific, |
| *ARRB1* | Arrestin Beta 1 | A_23_P203702 | 0.20* | 0.11* | 0.38 | Chan, Epiblast specific |
| ***ATG13***[b] | Autophagy Related 13 | A_23_P95292 | 1.20 | 1.05 | 1.37 | Tesar |
| ***CD44*** | CD44 Molecule (Indian Blood Group) | A_23_P24870 | 1.08 | 0.98 | 1.19 | High in Gafni PSCs |
| ***CD9*** | CD9 Molecule | A_23_P76364 | 0.94 | 0.88 | 1.00 | High in Takashima |
| ***CDH1*** | Cadherin 1 | A_23_P206359 | 1.58* | 1.38 | 1.82 | mESC specific |
| ***CLEC4D*** | C-Type Lectin Domain Family 4 Member D | A_23_P25235 | 1.09 | 1.14 | 1.04 | Epiblast specific in Chan 3iL PSCs |
| ***COL1A1*** | Collagen Type I Alpha 1 Chain | A_23_P207520 | 0.38* | 0.28 | 0.50 | High in Takashima & Chan PSCs |
| ***COMMD3*** | COMM Domain Containing 3 | A_23_P138514 | 0.63* | 0.60 | 0.67 | High in Takashima nPSCs |
| ***CTNNB1*** | Catenin Beta 1 | A_23_P29495 | 0.83 | 0.82 | 0.84 | High in Takashima nPSCs |
| ***DAZL*** | Deleted In Azoospermia Like | A_23_P212105 | 1.05 | 1.05 | 1.06 | mESC specific |
| ***DNMT3L*** | DNA Methyltransferase 3 Like | A_23_P17673 | 7.53* | 5.42 | 10.47 | Naive specific, Neri, Takashima, Theunissen |
| ***DPPA2*** | Developmental Pluripotency Associated 2 | A_23_P405885 | 1.69 | 2.03 | 1.41 | N list in Valamehr & Theunissen |
| ***DPPA5*** | Developmental Pluripotency Associated 5 | A_32_P233950 | 10.76* | 11.85 | 9.78 | List in Valamehr; Collier, Theunissen |
| ***DUSP6*** | Dual Specificity Phosphatase 6 | | 0.05* | 0.04* | 0.05* | |
| ***ESRRB*** | Estrogen Related Receptor Beta (not expressed in the human epiblast, High in t2iL+ dox inducible KLF2 and Nanog; Low in t2iL+ Gő) | A_24_P415928 | 1.01 | 1.03 | 1.00 | List in Valamehr & Takashima; Pera & Rossant |
| ***ETV5*** | ETS Variant Transcription Factor 5 | A_23_P9836 | 0.58* | 0.64 | 0.52 | High in Gafni PSCs |
| *FBXO15* | F-Box Protein 15 | A_23_P342709 | 4.67* | 6.13* | 3.55* | mESC specific |
| ***FGF4*** | Fibroblast Growth Factor 4 | A_24_P355720 | 3.18* | 2.55 | 3.95 | List in Valamehr, Takashima, & Chan |
| ***FGF8*** | Fibroblast Growth Factor 8 | A_23_P46829 | 0.69 | 0.70 | 0.69 | High in Takashima & Chan PSCs |
| ***FN1*** | Fibronectin 1 **Transcriptionally regulated with naive protocols** | A_24_P119745 | 0.36* | 0.19* | 0.68 | High in Tesar Takashima reports |
| ***GBX2*** | Gastrulation Brain Homeobox 2 | A_23_P131183 | 0.97 | 0.94 | 1.00 | mESC specific |
| ***GDF3*** | Growth Differentiation Factor 3 | A_23_P72817 | 0.98 | 0.85 | 1.14 | mESC specific, Chan |
| ***HORMAD1*** | HORMA Domain Containing 1 | A_32_P199884 | 1.26 | 1.44 | 1.10 | High in Tesar mESCs; Theunissen |
| ***ID3*** | Inhibitor of DNA Binding 3, HLH Protein | A_23_P137381 | 0.97 | 0.77 | 1.22 | High in Gafni PSCs |
| *ID4* | Inhibitor Of DNA Binding 4, HLH Protein | A_23_P59375 | 1.05 | 1.08 | 1.03 | List in Valamehr |
| ***IGFBP2*** | Insulin Like Growth Factor Binding Protein 2 | A_23_P119943 | 0.97 | 1.25 | 0.75 | High in Takashima nPSCs |
| ***IL6ST*** | Interleukin 6 Cytokine Family Signal Transducer, | A_32_P140656 | 0.93 | 0.73 | 1.19 | Epiblast specific CD130 (IL6ST, LIF-coreceptor) expressed in human epiblast, Collier) |
| ***KIT*** | KIT Proto-Oncogene, Receptor Tyrosine Kinase | A_23_P110253 | 1.18 | 0.99 | 1.42* | Takashima |

| | | | | | | |
|---|---|---|---|---|---|---|
| ***KLF2*** | Kruppel Like Factor 2<br>Not expressed in human epiblast | A_23_P119196 | **3.66*** | 2.51 | **5.33*** | In mESCs, Smith, Tesar, Takashima |
| ***KLF4*** | Kruppel Like Factor 4 | A_23_P32233 | **3.98*** | **3.95*** | **4.01*** | Naive specific marker, Tesar & Theunissen |
| ***KLF5*** | Kruppel Like Factor 5 | A_23_P53891 | **3.50*** | 2.52 | 4.86 | |
| ***LAMA1*** | Laminin Subunit Alpha 1 | A_24_P100613 | 0.74 | 0.59 | 0.92 | Takashima |
| ***LIFR*** | LIF Receptor Subunit Alpha | A_24_P397386 | **0.51*** | **0.35*** | 0.74 | Takashima |
| ***LIMCH1*** | LIM and Calponin Homology Domains 1 | A_32_P117354 | **2.07*** | 1.95 | 2.20 | Epiblast specific |
| ***MAEL*** | Maelstrom Spermatogenic Transposon Silencer | A_23_P114934 | 1.25 | 1.21 | 1.28 | Theunissen |
| *MFAP3L* | Microfibril Associated Protein 3 Like | A_24_P76675 | 0.85 | 0.72 | 1.00 | Epiblast specific |
| *MICA* | MHC Class I Polypeptide-Related Sequence A | A_23_P257516 | **1.47*** | 1.40 | **1.54*** | |
| *MICB* | MHC Class I Polypeptide-Related Sequence B | A_23_P387471 | **2.14*** | 2.08 | 2.20 | |
| ***NANOG*** | Nanog Homeobox<br>**Transcriptionally regulated with naive protocols** | A_23_P204640 | **0.64*** | 0.78 | 0.53 | Collier & Theunissen |
| ***NR0B1*** | Nuclear Receptor Subfamily 0 Group B Member 1 | A_23_P73632 | **0.38*** | **0.28*** | 0.51 | Epiblast specific, Tesar |
| *PIWIL2* | Piwi Like RNA-Mediated Gene Silencing 2 | A_32_P208654 | 0.58 | 0.86 | 0.39 | Epiblast specific |
| ***POU5F1*** | **Transcriptionally regulated with naive protocols** | A_24_P214841 | 0.74 | 0.64 | 0.85 | Theunissen |
| *PRDM14* | PR/SET Domain 14 | A_23_P123488 | 1.00 | 1.18 | 0.85 | Maintenance of pluripotency, Takashima |
| ***PYGL*** | Glycogen Phosphorylase L<br>**Transcriptionally regulated with naive protocols,** low in naïve mESCs | A_23_P48676 | 0.65 | 0.67 | 0.64 | Chan, Tesar |
| ***REST*** | **Transcriptionally regulated with naive protocols** | | **1.94*** | 1.90 | 1.99 | Takashima |
| *SLC25A16* | Solute Carrier Family 25 Member 16 | A_23_P423891 | **3.04*** | **3.11*** | **2.98*** | Epiblast specific |
| *SMYD2* | SET and MYND Domain Containing 2 | A_23_P170587 | 1.38 | 1.07 | 1.77 | Epiblast specific |
| *SOAT1* | Sterol O-Acyltransferase 1 | A_32_P103131 | **2.78*** | 2.11 | 3.65 | Epiblast specific |
| ***SOX2*** | SRY-Box Transcription Factor 2 | A_24_P379969 | 1.01 | 0.88 | 1.16 | mESC specific, Takashima |
| *STAT3* | Signal Transducer and Activator of Transcription 3 | A_23_P107206 | **1.94*** | 1.55 | **2.44*** | List in Valamehr |
| ***TDGF1*** | Teratocarcinoma-Derived Growth Factor 1 (Known as Cripto-1) | A_23_P366376 | 0.77 | 0.86 | 0.68 | Takashima |
| ***TERT*** | Telomerase Reverse Transcriptase | A_23_P110851 | **0.59*** | **0.69*** | **0.51*** | Takashima |
| ***TFCP2L1*** | Transcription Factor CP2 Like 1 | A_23_P5301 | **2.95*** | 3.66 | 2.37 | List in Valamehr; Takashima, Theunissen |
| ***TFE3*** | Transcription Factor Binding to IGHM Enhancer 3 | A_23_P84952 | 1.11 | 0.88 | 1.40 | Takashima |
| ***UTF1*** | Undifferentiated Embryonic Cell Transcription Factor 1 | A_23_P33141 | 1.43 | 1.50 | 1.36 | Llist in Valamehr; mESC specific, Theunissen |
| ***XIST*** | X Inactive Specific Transcript | A_24_P500584 | 1.09 | 1.11 | 1.06 | P list in Valamehr |
| ***ZFP42*** | ZFP42 Zinc Finger Protein, known as Rex1, a definitive naïve marker in the mouse | A_23_P395582 | 0.83 | 0.87 | 0.79 | High in Takashima, Theunissen; Low in Gafni, Chan, & Ware PSCs |
| *ZNF600* | Zinc Finger Protein 600 | A_23_P16006 | 1.22 | 1.01 | 1.48 | Epiblast specific |
| **Formative (between pre-implantation and the E6.5 epiblast in mouse) gene expression** (n = 5, * indicates $P < 0.05$) | | | | | | |
| ***DNMT3A*** | DNA Methyltransferase 3 Alpha<br>**Transcriptionally regulated with naïve protocols, decrease in naïve** | A_23_P154500 | 0.98 | 0.97 | 0.98 | Smith review, Gafni |
| ***DNMT3B*** | DNA Methyltransferase 3 Beta<br>**Transcriptionally regulated with naïve protocols, decrease in naive** | A_23_P28953 | 0.99 | 0.96 | 1.02 | Smith review; Decreased in Takashima, Gafni, & Neri |
| ***OTX2*** | Orthodenticle Homeobox 2<br>**Transcriptionally regulated with naive protocols** | A_23_P48663 | **0.52*** | 0.52 | 0.52 | Formative state, priming marker (Smith); Chan, Gafni, Theunissen, Valamehr, Ware |
| ***SALL2*** | Spalt Like Transcription Factor 2 | A_23_P48585 | **0.74*** | **0.78*** | 0.59 | Smith review |
| ***SOX3*** | SRY-Box Transcription Factor 3 | A_23_P85218 | 0.60 | 0.82 | 0.44 | Smith review |
| **Primed or metastable lineage specifier gene expression** (n = 37, * indicates $P < 0.05$) | | | | | | |
| *ACTC1* | Actin Alpha Cardiac Muscle 1 | A_23_P205894 | **0.09*** | **0.10*** | **0.08*** | |

| | | | | | | |
|---|---|---|---|---|---|---|
| *BMP2* | Bone Morphogenetic Protein 2 | A_23_P143331 | **0.28*** | 0.20 | 0.40 | |
| *CD47* | CD47 Molecule | A_23_P6935 | 0.77 | 0.82 | 0.72 | |
| *CDH11* | Cadherin 11 | A_23_P152305 | 1.33 | 1.54 | 1.15 | P list in Valamehr |
| ***CER1*** | Cerberus 1, DAN Family BMP Antagonist | A_23_P329798 | **0.27*** | 0.34 | **0.22*** | High in EpiSCs (Tesar & Brons); low in Chan & Takashima PSCs |
| *COL13A1* | Collagen Type XIII Alpha 1 Chain | A_23_P1331 | **0.28*** | 0.27 | **0.28*** | P list in Valamehr |
| *CXCR4* | C-X-C Motif Chemokine Receptor 4 | A_23_P102000 | **0.19*** | **0.15*** | 0.25 | P list in Valamehr |
| *CYP2B6* | Cytochrome P450 Family 2 Subfamily B Member 6 | A_24_P339514 | **0.32*** | 0.40 | 0.25 | P list in Valamehr |
| *DLL1* | Delta Like Canonical Notch Ligand 1 | A_23_P167920 | **2.86*** | 2.29 | 3.57 | |
| ***DNMT1*** | DNA Methyltransferase 1 | A_24_P408083 | 0.90 | 0.80 | 1.01 | Zhao |
| ***EGR1*** | Early Growth Response 1 | A_23_P214080 | **0.01*** | **0.01*** | **0.01*** | P list in Valamehr |
| *EOMES* | Eomesodermin | A_24_P97374 | **0.18*** | 0.09 | 0.36 | |
| *ERBB4* | Erb-B2 Receptor Tyrosine Kinase 4 | A_32_P183765 | **0.08*** | **0.07*** | **0.09*** | P list in Valamehr |
| *FLT1* | Fms Related Receptor Tyrosine Kinase 1 | A_24_P576191 | **0.22*** | 0.34 | 0.15 | |
| *FOXA2* | Forkhead Box A2 | A_24_P365515 | **0.15*** | **0.07*** | 0.36 | |
| *GAL* | Galanin And GMAP Prepropeptide | A_23_P374844 | **0.22*** | **0.18*** | 0.27 | P list in Valamehr |
| *GSC* | Goosecoid Homeobox | A_23_P76774 | **0.07*** | **0.06*** | **0.08*** | P list in Valamehr |
| *HEPH* | Hephaestin | A_24_P399980 | **0.74*** | 0.73 | 0.74 | P list in Valamehr |
| *HES1* | Hes Family BHLH Transcription Factor 1 | A_23_P17998 | **0.26*** | 0.31 | 0.22 | P list in Valamehr |
| *HHEX* | Hematopoietically Expressed Homeobox | A_23_P47034 | 0.70 | 0.50 | 0.97 | Pera & Rossant |
| ***LEFTY1*** | Left-Right Determination Factor 1 | A_23_P160336 | **0.04*** | 0.08 | 0.02 | P list in Valamehr Mallon, Chen |
| ***LEFTY2*** | Left-Right Determination Factor 2 | A_23_P137573 | **0.18*** | 0.27 | 0.13 | Mallon, Chen |
| *MEGF10* | Multiple EGF Like Domains 10 | A_24_P7192 | 1.74 | 2.15 | 1.41 | P list in Valamehr |
| ***MEIS1*** | Meis Homeobox 1 **Transcriptionally regulated with naive protocols** | A_24_P319736 | 1.24 | 1.55 | 1.00 | Low in Gafni PSCs |
| ***MEIS2*** | Meis Homeobox 2 **Transcriptionally regulated with naive protocols** | A_24_P82200 | 1.59 | 1.62 | 1.56 | Low in Gafni PSCs |
| *NIPBL* | NIPBL Cohesin Loading Factor | A_24_P180383 | 0.83 | 0.81 | 0.85 | P list in Valamehr |
| ***NODAL*** | **Transcriptionally regulated with naive protocols** | A_23_P127322 | 0.48 | 0.59 | 0.38 | Pera & Rossant |
| *NR2F2* | Nuclear Receptor Subfamily 2 Group F Member 2 | A_23_P88589 | 1.33 | 1.76 | 1.00 | P list in Valamehr |
| *PCDH10* | Protocadherin 10 | A_32_P168605 | 0.79 | 0.63 | 1.00 | P list in Valamehr |
| ***PITX2*** | Paired Like Homeodomain 2 | A_23_P167367 | **0.06*** | 0.08 | **0.05*** | P list in Valamehr |
| *RORA* | RAR Related Orphan Receptor A | A_23_P26124 | 0.58 | 0.70 | 0.48 | P list in Valamehr |
| *SOX1* | SRY-Box Transcription Factor 1 | A_24_P112803 | 1.37 | 1.37 | 1.37 | Pera & Rossant |
| *SOX11* | SRY-Box Transcription Factor 11 associated with neuron differentiation | A_24_P302584 | 1.30 | 1.40 | 1.21 | Low in Gafni PSCs |
| *TGFB2* | Transforming Growth Factor Beta 2 | A_24_P148261 | **0.69*** | 0.82 | 0.58 | |
| *TOX* | Thymocyte Selection Associated High Mobility Group Box | A_24_P226755 | 1.21 | 1.33 | 1.10 | P list in Valamehr |
| *ZIC1* | Zic Family Member 1 | A_23_P367618 | 1.00 | 1.00 | 1.00 | Low in Gafni PSCs |
| *ZIC2* | Zic Family Member 2 | A_23_P36972 | 0.83 | 0.89 | 0.77 | Priming marker (Smith) |
| **Other X-lined or imprinted gene expression** (n = 5, * indicates $P < 0.05$) | | | | | | |
| ***DLK1*** | PEG9, Paternally Expressed 10, Known as DLK1, Delta Like Non-Canonical Notch Ligand 1 | A_24_P236251 | 3.41 | 2.56 | 4.54 | Low in naive & high in primed states in Gafni |
| *FMR1* | FMRP Translational Regulator 1, X-linked | A_24_P93967 | 1.30 | 1.80 | 0.93 | |
| ***H19*** | H19, Imprinted Maternally Expressed | A_24_P52697 | 1.60 | 0.90 | 2.87 | |
| ***MEG3*** | MEG3, Paternally Imprinted gene, Known as GTL2 **Transcriptionally regulated with naive protocols** | A_24_P272993 | 1.52 | 1.25 | 1.85 | High in naïve (Collier) |
| ***MEST*** | MEST, Mesoderm specific, imprinted, preferentially paternally expressed | A_23_P156970 | 0.92 | 0.92 | 0.93 | |

### FOOTNOTES

[a] Based on the references as detailed below.

**[b]** Transcriptionally regulated genes are indicated in bold gene symbols, with up-regrated gene in red and down-regulated gene in blue colors.

* Indicates *P* values < 0.05 with two-tailed Student *t*-test, for fold changes of gene expression in microarray.

Additional abbreviations: EPS, epiblast specific gene signature; mESCs, naive mouse embryonic stem cells; N, naive-related genes; nPSCs, naive pluripotent stem cells; PSCs, pluripotent stem cells; P, primed or primed genes or cells.
