## Supplementary material for "Resistance to Naïve and Formative Pluripotency Conversion in RSeT Human Embryonic Stem Cells": Key Resource Table

**KEY RESOURCES TABLE**

| REAGENT or RESOURCE | SOURCE | IDENTIFIER |
|---|---|---|
| **Chemicals** | | |
| Dimethyl sulfoxide (DMSO) | Sigma | D2650 |
| Dorsomorphin | Sigma | P5499 |
| Erlotinib HCl (OSI-744) | Selleck Chemicals | S1023 |
| Gö6983 | Sigma | G1918-1MG |
| Gö6983 | Sigma | G1918 |
| JAK inhibitor I | EMD4 Biosciences | 420099 |
| Paraformaldehyde (PFA) | Electron Microscopy Sciences | 15710 |
| PD173074 | Selleck Chemicals | S1264 |
| SB431542 | Tocris Bioscience | 1614 |
| Y-27632 | EMD4 Biosciences | 688000 |
| Y-27632 dihydrochloride | Tocris Bioscience | 1254 |
| β-mercaptoethanol (14.3 M) | Sigma | M6250 |
| **Antibodies** | | |
| BD Pharmingen™ Alexa Fluor® 647 Mouse Anti-Human CD130; Mouse BALB/c IgG1, κ; Clone: AM64 | BD Biosciences | 564151 |
| BD Pharmingen™ Alexa Fluor® 647 Mouse Anti-Human CD75/CD75s, IgM, κ, Clone: ZB55 | BD Biosciences | 566350 |
| BD Pharmingen™ Alexa Fluor® 647 Mouse Anti-Human SUSD2, IgG1, κ, Clone: W5C5 | BD Biosciences | 566657 |
| BD Pharmingen™ Alexa Fluor® 647 Mouse IgM, κ Isotype Control, Clone: G155-228 | BD Biosciences | 560806 |
| BD Pharmingen™ FITC Mouse Anti-Human CD57 Mouse IgM, κ, Clone: NK-1 | BD Biosciences | 555619 |
| BD Pharmingen™ FITC Mouse IgG2a, κ Isotype Control, Clone: G155-178 | BD Biosciences | 555573 |
| BD Pharmingen™ FITC Mouse IgM, κ Isotype Control, Clone: G155-228 | BD Biosciences | 551448 |
| BD Phosflow™ Alexa Fluor® 647 Mouse IgG1 κ Isotype control, MOPC-21 | BD Biosciences | 557783 |
| BD™ CD24 FITC, Isotype: Mouse IgG2a, κ, Clone: ML5 | BD Biosciences | 655154 |
| Goat anti-Mouse IgG1 Secondary Antibody, Alexa Fluor® 488 conjugate | Thermo Fisher Scientific | A-21121 |

| | | |
|---|---|---|
| Goat anti-Mouse IgG1 Secondary Antibody, Alexa Fluor® 647 conjugate | Thermo Fisher Scientific | A-21240 |
| Goat anti-Mouse IgG2a Secondary Antibody, Alexa Fluor® 555 conjugate | Thermo Fisher Scientific | A-21137 |
| Goat anti-Mouse IgG2a Secondary Antibody, Alexa Fluor® 647 conjugate | Thermo Fisher Scientific | A-21241 |
| Goat anti-Mouse IgG2b Secondary Antibody, Alexa Fluor® 647 conjugate | Thermo Fisher Scientific | A-21242 |
| Goat anti-Mouse IgM Heavy Chain Secondary Antibody, Alexa Fluor® 647 conjugate | Thermo Fisher Scientific | A-21238 |
| Goat anti-Mouse IgM Heavy Chain Secondary Antibody, Alexa Fluor® 647 conjugate | Thermo Fisher Scientific | A-21238 |
| NANOG (rabbit IgG) | ReproCELL Inc, Japan | RCAB0004P-F |
| Oct-4, mouse IgG2b | Santa Cruz Biotechnology | sc-5279 |
| Pharmingen™ FITC Mouse Anti-Human CD90, Isotype: Mouse IgM, κ; Clone: 5E10 | BD Biosciences | 561969 |
| SSEA-1, mouse IgM | Santa Cruz Biotechnology | sc-21702 |
| SSEA-4, mouse IgG3 | Santa Cruz Biotechnology | sc-21704 |
| Tra-1-60, mouse IgM | Santa Cruz Biotechnology | sc-21705 |
| Tra-1-81, mouse IgM | Santa Cruz Biotechnology | sc-21706 |
| **Cell Culture Reagents** | | |
| Accutase™ | Innovative Cell Technologies | AT-104 |
| BD Falcon Cell Strainer | BD Bioscience | 352340 |
| Bovine Albumin Fraction V (7.5% solution) | Thermo Fisher Scientific | 15260037 |
| CryoStor CS10 | StemCell Technologies | 7930 |
| DMEM/F12, no HEPES | Thermo Fisher Scientific | 11320-082 |
| DMEM/F12, with HEPES | Thermo Fisher Scientific | 11330–032 |
| Dulbecco's Phosphate-Buffered Saline | Thermo Fisher Scientific | 14190-144 |
| Faxitron Cabinet X-ray System | Faxitron X-ray Corporation, Wheeling, IL | Model RX-650 |
| Fetal Bovine Serum (FBS), certified, heat inactivated | Thermo Fisher Scientific | 10082147 |
| Heat-inactivated fetal bovine serum (FBS) | Hyclone (Logan Utah) | SH30071-03 |
| hESC-qualified Matrigel | BD Bioscience | 354277 |
| Knockout Serum Replacer | Thermo Fisher Scientific | 10828–028 |
| L-Glutamine (200 mM) | Thermo Fisher Scientific | 25030–081 |
| MEM non-essential amino acids solution (100X) | Thermo Fisher Scientific | 11140050 |
| mTeSR1 and Supplements | StemCell Technologies | 5850 |
| MULTIWELL six-well plates | Becton Dickinson Labware | 353046 |

| | | |
|---|---|---|
| Nalgene 5100-0001 Cryo 1°C | Thermo Fisher Scientific | C6516F-1 |
| RSeT™ Medium (2-Component) | StemCell Technologies | 05978 |
| Thermo Scientific Nunc Thermanox Coverslips (25 mm diameter; 500/cs) | Thermo Fisher Scientific | 174985 (25 mm diameter) |
| TrypLE™ Express | Thermo Fisher Scientific | REF 12604-013 |
| Trypsin | Thermo Fisher Scientific | 25300-054 |
| **Critical Commercial Assays** | | |
| Countess™ automated cell counter (assay) | Thermo Fisher Scientific | C10227 |
| **Deposited Data** | | |
| Microarray | Agilent Technologies, Inc. | In this study |
| **Experimental Models: Cell Lines** | | |
| H1 (WA01) | WiCell Inc. | NIHhESC-10-0043 |
| H7 (WA07) | WiCell Inc. | NIHhESC-10-0061 |
| H9 (WA09) | WiCell Inc. | NIHhESC-10-0062 |
| **Real-time PCR Reagents** | | |
| Nuclease-free water | Thermo Fisher Scientific | R0582 |
| SuperScript® VILO™ cDNA Synthesis Kit | Thermo Fisher Scientific | 11754050 |
| TagMan Assay ID Hs99999903_m1 for *ACTB* | Thermo Fisher Scientific | 4331182 |
| TagMan Assay ID Hs99999905_m1 for *GAPDH* | Thermo Fisher Scientific | 4453320 |
| TaqMan® Fast Advanced Master Mix | Thermo Fisher Scientific | 4444557 |
| TE, pH 8.0, RNase-free | Thermo Fisher Scientific | AM9849 |
| **Recombinant DNA** | | |
| None | | |
| **Software and Algorithms** | | |
| FlowJo | BD Biosciences | Version 10.9.0 |
| ImageJ | NIH, Bethesda, USA | |
| QuantStudio™ 6 and 7 Flex Real-Time PCR System Software | Thermo Fisher Scientific | Publication Number 4489822 |
| R software package | http://cran.r-project.org/ | |
| SC3 consensus clustering | Bioconductor (http://bioconductor.org) | Version 3.12 |
